# Integrated Drivers of Basal and Acute Immunity in Diverse Human Populations

**DOI:** 10.1101/2023.03.25.534227

**Authors:** Aisha Souquette, E. Kaitlynn Allen, Christine M. Oshansky, Li Tang, Sook-san Wong, Trushar Jeevan, Lei Shi, Stanley Pounds, George Elias, Guillermina Kuan, Angel Balmaseda, Raul Zapata, Kathryn Shaw-Saliba, Pierre Van Damme, Viggo Van Tendeloo, Juan Carlos Dib, Benson Ogunjimi, Richard Webby, Stacey Schultz-Cherry, Andrew Pekosz, Richard Rothman, Aubree Gordon, Paul G. Thomas

**Affiliations:** Department of Immunology, St. Jude Children’s Research Hospital, Memphis, Tennessee, 38105, USA; University of Tennessee Health Science Center, Memphis, Tennessee, 38163, USA; Department of Biostatistics, St. Jude Children’s Research Hospital, Memphis, Tennessee, 38105, USA; Department of Infectious Disease, St. Jude Children’s Research Hospital, Memphis, Tennessee, 38105, USA; Laboratory of Experimental Hematology (LEH), Vaccine and Infectious Disease Institute, University of Antwerp, Antwerp, 2610, Belgium; Center for Health Economics Research and Modeling Infectious Diseases (CHERMID), Vaccine and Infectious Disease Institute, University of Antwerp, Antwerp, 2610, Belgium; Antwerp Center for Translational Immunology and Virology (ACTIV), Vaccine and Infectious Disease Institute, University of Antwerp, Antwerp, 2610, Belgium; Antwerp Unit for Data Analysis and Computation in Immunology and Sequencing (AUDACIS), University of Antwerp, Antwerp, 2610, Belgium; Centro de Salud Sócrates Flores Vivas, Ministry of Health, Managua, 12014, Nicaragua; Centro Nacional de Diagnóstico y Referencia, Ministry of Health, Managua, 16064, Nicaragua; Sustainable Sciences Institute (Site - Managua, Nicaragua), San Francisco, California, 94102, USA; Department of Emergency Medicine, Johns Hopkins University School of Medicine, Baltimore, MD, 21209, USA; Center for the Evaluation of Vaccination (CEV), Vaccine and Infectious Disease Institute, University of Antwerp, Antwerp, 2610, Belgium; Department of Medicine, Universidad del Norte, Barranquilla, 081001, Colombia; Department of Paediatrics, Antwerp University Hospital, Antwerp, 2610, Belgium; W. Harry Feinstone Department of Molecular Microbiology and Immunology, The Johns Hopkins Bloomberg School of Public Health, Baltimore, Maryland 21205, USA; Department of Epidemiology, School of Public Health, University of Michigan, Ann Arbor, Michigan, 48109, USA

**Keywords:** immune variation, influenza, herpesvirus, co-infection, systems immunology, basal immunity, acute immunity, immunogenetics, immunophenotype, correlate of severity

## Abstract

Prior studies have identified genetic, infectious, and biological associations with immune competence and disease severity; however, there have been few integrative analyses of these factors and study populations are often limited in demographic diversity. Utilizing samples from 1,705 individuals in 5 countries, we examined putative determinants of immunity, including: single nucleotide polymorphisms, ancestry informative markers, herpesvirus status, age, and sex. In healthy subjects, we found significant differences in cytokine levels, leukocyte phenotypes, and gene expression. Transcriptional responses also varied by cohort, and the most significant determinant was ancestry. In influenza infected subjects, we found two disease severity immunophenotypes, largely driven by age. Additionally, cytokine regression models show each determinant differentially contributes to acute immune variation, with unique and interactive, location-specific herpesvirus effects. These results provide novel insight into the scope of immune heterogeneity across diverse populations, the integrative effects of factors which drive it, and the consequences for illness outcomes.

## Introduction

Functional consequences of immune heterogeneity are demonstrated by variation in vaccine efficacy, responses to therapeutic treatment, infectious disease severity, and susceptibility to autoimmunity. Understanding the landscape and drivers of immune variation would provide valuable insight into immune competence and points of intervention to improve disease outcome. Indeed, baseline immune predictors of vaccine responses and neoplastic disease prognosis have already been identified in a variety of studies.^1,2^

Immune variation, at baseline and in response to immune challenge, is dynamically influenced by unique and interactive effects of intrinsic and extrinsic host factors. Intrinsic factors are internal to the host and include variables such as age, sex, genetics, and microbiome. Extrinsic factors are external to the host and include variables such as diet, environment, and acquired infections. Each of these determinants can influence multiple facets of immunity, including, but not limited to, susceptibility to infection or immune mediated disease, the basal activation state of the immune system, pathogen detection, downstream signaling, priming of adaptive immunity, leukocyte phenotype and function, and/or response to therapeutic treatment.^3–22^ For example, in a study of three independent influenza cohorts, we found that a single nucleotide polymorphism rs3448144-A in *IFITM3* is enriched in severe patients, governs transcription factor binding and promoter activity, and alters methylation of the *IFITM3* promoter in CD8 T cells, which correlates with cell frequency in nasal washes.^12^ Coinciding with these results, recent studies have reported associations between rs3448144 and COVID-19 disease severity.^23,24^ Importantly, the collective impact of immune modulation induced by intrinsic and extrinsic host factors can alter vaccination responses and illness outcome. However, further study is needed to clarify the quantitative effects of these variables, as conflicting conclusions have occurred across studies, possibly due to skews in age distribution, geographical location, or ancestry impacting observed results. Ultimately, this limits both our understanding of the extent of immune heterogeneity and the general applicability of conclusions drawn from single cohorts.

Here, we quantify the contribution of intrinsic and extrinsic host factors on the setpoint of baseline immunity, the transcriptional response to *in vitro* immune challenge, and the magnitude and quality of the immune response to *in vivo* influenza infection in diverse populations around the world. Immune measures include cytokines (from plasma and nasal wash), cellular phenotypes, and/or gene expression (both at baseline and post PBMC stimulation with PRR ligands). Of these, cytokines were our primary and most extensively used measurement, with data collected at the RNA and protein level. While most prior work has focused on cell phenotypes (by flow cytometry), cytokines have several advantages from both a scientific and clinical standpoint. Samples are easily obtained and stored, and analytes can quickly and precisely be measured with minimal equipment, sample input, and processing. In addition, they have a substantial dynamic range and offer broad profiling of innate and adaptive immunity. For influenza infected subjects, outcome measures include presence/absence of symptoms, symptom severity scores, viral load, and duration of influenza virus shedding. To explain variation in these measures, linear regression modeling was utilized, which allows for an easily interpretable and transparent integrative analysis of all independent variables and a quantification of their unique, interactive, and collective effects on the immune measure of interest. Collectively, the data presented here, representing 123,178,117 data points from 1,705 individuals, show age, sex, herpesviruses, and host genetics can independently and interactively influence baseline immunity and the acute immune response to influenza virus infection. Of these determinants, ancestry is the most significant factor. Importantly, these factors differentially contribute to variation in a given immune measure, and the cumulative impact of immune modulation by these factors results in unique immune profiles associated with disease severity. Our results demonstrate that simultaneous assessment of diverse populations is necessary to better understand the scope of immune variation and the factors that contribute to it.

## Results

### Explanatory factor variation within and across diverse populations

This study includes samples from 1,705 healthy or influenza infected subjects from 5 countries and 8 distinct populations (**Fig 1**). North American populations include the Memphis cohort and the Baltimore cohort. Latin American populations include the Nicaragua cohort and the Colombia cohorts, which include subjects from the local city of Calabazo and two indigenous populations, Seywiaka and Umandita. Samples also include a European cohort from Belgium and an Asian cohort from Taiwan. Select assays and corresponding analyses focused on our two most diverse cohorts, Memphis and Nicaragua, which have the largest sample size, availability, and variety of sample types collected. These populations also capture environmental diversity, and, collectively, represent three distinct racial/ethnic groups (White/Caucasian, Hispanic/Latino, Black/African American), two of which are highly admixed, thereby representing greater genetic diversity. Enrichment of diverse, understudied racial/ethnic groups is a key feature of our study, as recent work has shown that increasing genetic diversity, as opposed to increasing the sample size of European ancestry groups, provides substantial improvement to genetic analyses, such as fine-mapping functional variants and portability of polygenic risk scores.^25,26^

**Figure 1.**
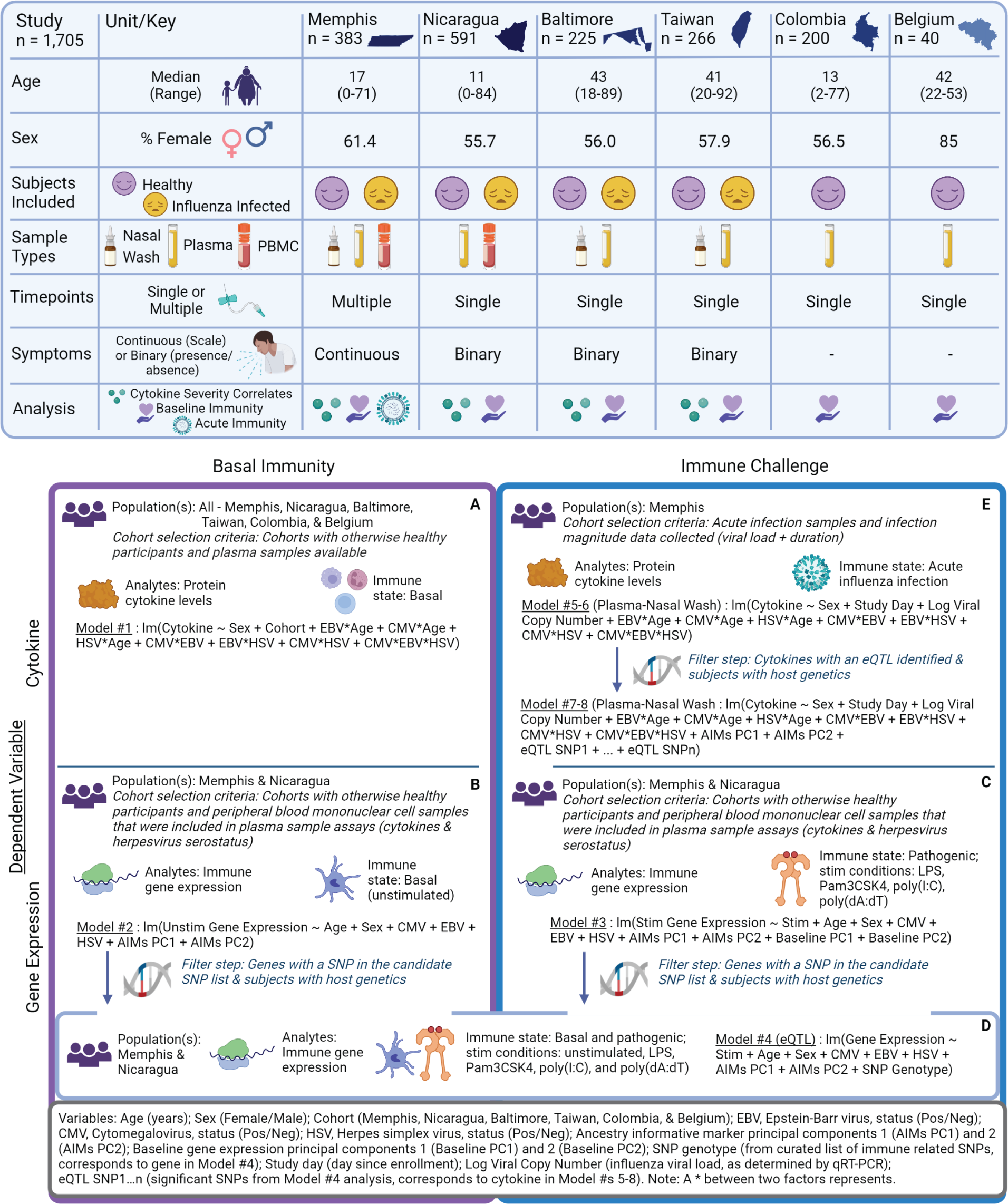
Study populations and model summary. Table of cohort characteristics (top) and overview of regression model analyses (bottom, labeled A-D in order of appearance).

Before examining baseline and acute immunity, we assessed the degree of variation in explanatory variables across cohorts. Here, intrinsic host factors include biological and genetic variables. Biological factors include age and sex, and the distribution of each varied by cohort (**Fig 2A-B**). Overall, the study was 58% female and age ranged from <1 year up to 92 years of age.

**Figure 2.**
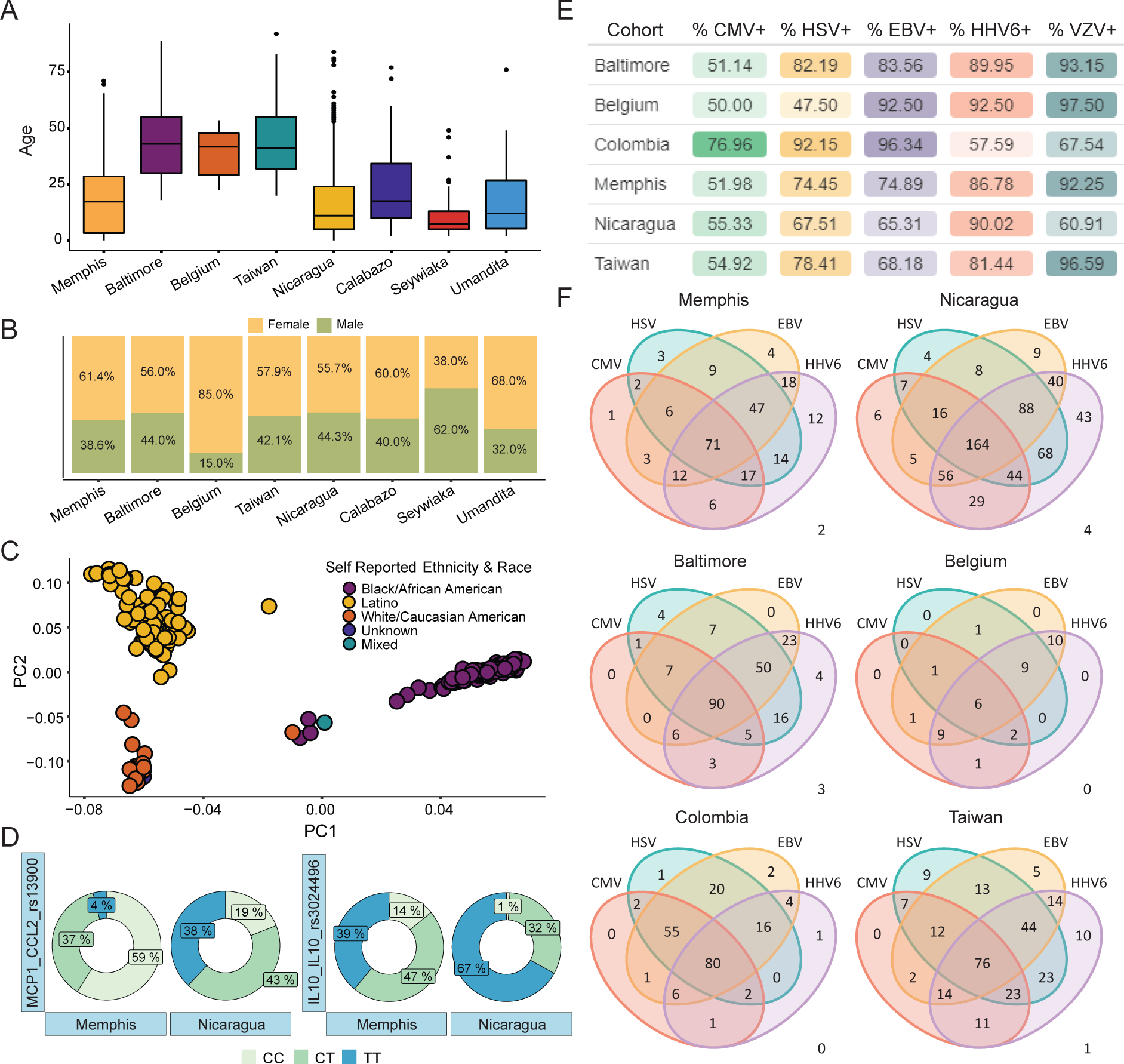
Variation of explanatory factors across diverse populations. Age (A) and sex (B) distribution for each cohort (n = 1,705). (C) Results from the principal component analysis of AIMs (n = 285). (D) Donut plots show the proportion of individuals in each cohort with a given genotype for a SNP of interest (n = 352), plot labels are encoded as follows: Protein-Gene-SNP. (E) Table of herpesvirus serostatus (%) by cohort (n = 1,532). (F) Venn-diagram of herpesvirus infection by cohort. Numbers in the bottom right represent the number of individuals negative for all indicated herpesviruses. For e sier visualization, the herpesvirus with the highest overall seroprevalence (VZV) was excluded.

Heritable contribution to immune variation was determined for a subset of healthy and infected subjects with available DNA or PBMCs (n = 355, 234 Memphians and 121 Nicaraguans), using genome-wide ancestry informative markers and candidate functional, immune-related SNPs. Genetic based ancestry, as determined by ancestry informative markers (AIMs), accounts for inherent genetic differences across diverse populations, typically from distinct geographical regions. This allows for a genetic and quantitative representation of admixed populations, as opposed to discrete classifications of race, which do not adequately represent populations with mixed ancestral backgrounds. AIMs were analyzed utilizing a principal component analysis (PCA), and the top two principal component values were included in downstream regression analyses. Graphical representation of the PCA results shows distinct clusters that were similar to the self-reported racial and ethnic makeup of the Memphis (75% African/Black American and 25% Caucasian/White American) and Nicaragua (100% Latino) cohorts, with a subset of ancestrally mixed individuals falling between clusters (**Fig 2C**). Importantly, only AIMs were able to distinguish between ancestral population clusters (**Supp Fig 1A-B**) and account for genetic variation within a given population (**Supp Fig 1C**). This highlights the limited applicability of current “Race” and “Ethnicity” categories collected in the clinical setting and applied in biomedical research, and supports current arguments for the need for inclusion of more appropriate factors in human studies aiming to distinguish between biological, epidemiological, and social determinants of health and disease.^27,28^ For the candidate SNP list, we examined common SNPs in functional regions of genes encoding the 41 cytokines in our assay and in 3 transcription factors important in initiating antiviral immunity, namely IRF3, IRF7, and all 3 subunits of NFkB (detailed description of prioritization is in the Methods). While some SNPs have similar allele frequencies across the Memphis and Nicaragua cohorts, several, such as rs13900 in MCP1 and rs3024496 in IL10, show significant variation and opposing major alleles in the two populations (**Fig 2D, Supp Table 1**), indicating genetic differences in immune-related genes with potential functional consequences due to ancestry.

Herpesviruses were utilized as an extrinsic factor of infectious exposure because they are globally prevalent, cause chronic infections that are generally well tolerated by immunocompetent hosts, and have previously been associated with modulation of the immune system.^29–32^ The serostatus of cytomegalovirus (CMV), Epstein-Barr virus (EBV), herpes simplex virus (HSV, 1 and 2 combined), human herpesvirus 6 (HHV6), and varicella-zoster virus (VZV) were determined for healthy and influenza infected subjects with available plasma from all eight cohorts (n = 1,532). Herpesvirus seroprevalence ranged from as low as 47.5% up to 97.5% (**Fig 2E**). Although the median age varies across cohorts, comparisons can be made between cohorts of similar age ranges and/or with age adjustment and these results demonstrate how herpesvirus prevalence varies across distinct groups. For example, although the Baltimore, Taiwan, and Belgium cohorts are similar in age, Belgium has 31-35% lower prevalence of HSV but a 9-24% higher prevalence of EBV. Additionally, despite having a similar age group, the Colombian cohort has 22% higher prevalence of CMV, 31% higher prevalence of EBV, and a 32% lower prevalence of HHV6 compared to Nicaragua. Importantly, with the exception of the Belgium cohort, which has the most limited age range in participants, each cohort exhibits a positive correlation between age and the total number of herpesviruses an individual has (**Supp Fig 1D**). This is particularly important to account for in downstream analyses because influenza disease severity is age associated, very young individuals (infants and children under 5) tend to have more severe infection and tend to be herpesvirus seronegative. Highlighting the ubiquity of herpesviruses, of the 1,532 individuals in this analysis, only 10 (0.587%) were negative for CMV, HSV, EBV, and HHV6, and only 118 (6.92%) tested positive for a single herpesvirus (**Fig 2F**). These data underscore the importance of accounting for multiple herpesvirus infections when trying to assess their contributions to a given disease outcome, as most individuals have more than one. However, due to the high prevalence of HHV6 and VZV, the herpesvirus serostatuses in downstream analyses are focused on CMV, EBV, and HSV.

### Distinct basal immune profiles mold response to immune challenge

To begin assessing baseline immune status, the levels of 38 plasma/serum cytokines were measured in 631 healthy subjects.^33–38^ Cytokine levels were natural log (ln) transformed and visualized using a t-Distributed Stochastic Neighbor Embedding (t-SNE) plot which aids in dimensionality reduction while utilizing data from all cytokines measured. These results show marked variation in cytokine levels, with distinct clustering by cohort. Some populations exhibit tight clusters (Baltimore), whereas others exhibit dispersed clusters (Colombia) or even two distinct clusters (Memphis), suggesting the variance of cytokine profiles across individuals varies by population and, consequently, focusing on a population with minimal variation across subjects may underestimate the extent of immune variation and bias results (**Fig 3A**). Split clustering of the Memphis cohort was not due to age, sex, herpesvirus serostatus, race, season/year, assay plate, or time between assay runs (**Supp Fig 2A-G**). Moreover, in a subsequent analysis, cohort clusters and their positions relative to each other remain after removal of Memphis cohort data (**Supp Fig 2H**). Variation was also observed at the individual cytokine level for immunomodulatory cytokines, pro-inflammatory cytokines, chemokines, and growth factors (**Fig 3B**).

**Figure 3.**
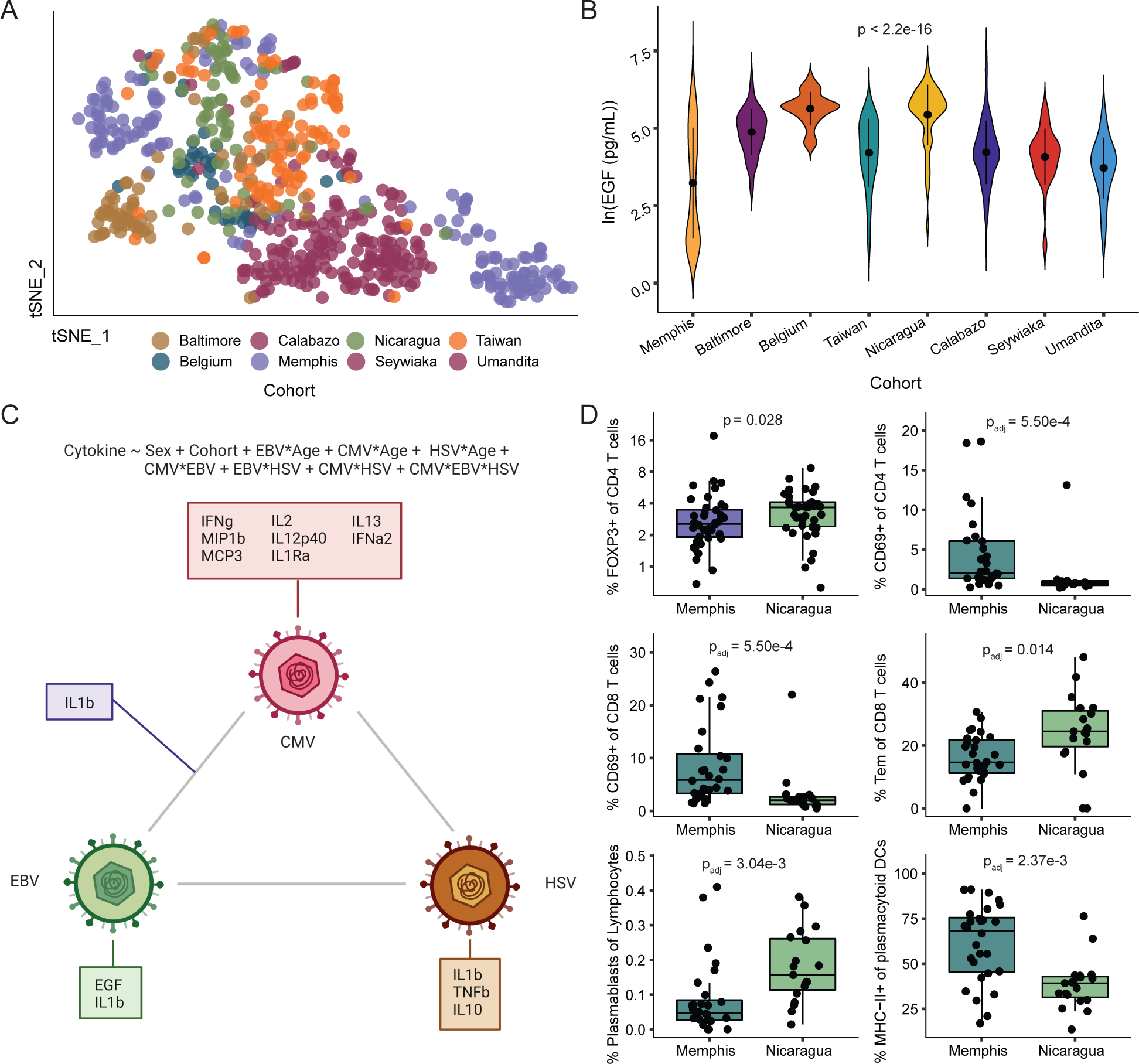
Basal immune profiles in ancestrally and environmentally distinct populations. (A) A t-SNE plot based on plasma/serum cytokines and colored by cohort (n = 631). (B) Violin plot demonstrates individual cytokine variation for EGF (n = 623). Black points in the center of the violin plot represent the mean cytokine level and line segments represent +/- the standard deviation. (C) Schematic of herpesvirus effects on basal cytokine levels based on linear regression modeling results (model displayed above diagram, n ≈ 500). Cytokines listed have an adjusted R^2^ value > 0.08 and FDR < 0.05. (D) Boxplots depict differences in basal cell phenotypes in PBMC samples from healthy subjects in the Memphis (n = 28) and Nicaragua (n = 19) cohorts. Presented p-values are from Mann-Whitney U tests, post adjustment for multiple comparisons (when applicable).

To examine the effects of intrinsic and extrinsic host factors on basal cytokine levels, a linear regression model (Model #1, **Fig 1**) was used to take into account the effects of: age, sex, cohort (accounting for genetics and unknown environmental factors specific to the geographically distinct populations), each herpesvirus alone, any interaction between a herpesvirus and age (due to the age association with serostatus), and any interaction between herpesviruses (since most individuals are infected with more than one). These results show herpesviruses have largely unique effects on basal cytokine levels, and the herpesvirus which caused the most effects was CMV (**Fig 3C** and **Supp Fig 3**). Each additional standard deviation unit increase in age (years) is cumulatively associated with altered GRO, IFNg, Eotaxin, and IL12p70, thus the larger the difference in age, the greater the magnitude of age effects (**Supp Fig 3**). Male sex was associated with decreased TGFa and cohort differentially affected all cytokines (**Supp Fig 3**). The adjusted R^2^ result from a linear regression model is an estimate of how much variability in the dependent variable the model can account for, and the “adjustment” refers to a penalty to the R^2^ value due to the incorporation of additional variables. This value was highest for the IL4 model at 0.4709 (47.1%) (**Supp Fig 3**). Another output from the regression models is an estimate of a coefficient for each factor which represents the change in the dependent variable per unit change in the factor. Comparing coefficients across factors provides insight into the magnitude of the effect of that factor. In these healthy cytokine models, cohort was associated with the strongest effects on circulating cytokine levels (**Supp Fig 3**). Collectively, these data suggest distinct populations have unique cytokine profiles, the effect of which likely represents divergent baseline immune states, significant impacts on downstream effector functions, including immune cell differentiation and functional responses to stimuli.

To further explore the potential for distinct immunophenotypes, we focused on two cohorts with PBMC samples, Memphis and Nicaragua, which allowed for more comprehensive investigation and statistical analysis. Flow cytometry was utilized to characterize the activation and/or differentiation state of circulating lymphocytes and antigen presenting cells at baseline. Samples included 28 Memphians (average age = 22, median age = 18) and 19 Nicaraguans (average age = 20, median age = 15). Nicaraguans exhibited a trend towards higher regulatory T cell (Treg)-like CD25+ CD4 T cells (padj = 0.071, **Supp Table 2**). To validate these findings, the Treg transcription factor FOXP3 (forkhead box protein P3) was included in a follow up experiment and results confirmed Nicaraguans have an increased frequency of FOXP3+ CD4 Tregs (p = 0.028, **Fig 3D**). Nicaraguans also showed significantly lower frequency and number of activated CD4 and CD8 T cells (padj = 5.5e-4 for both subsets), but higher plasmablasts (padj = 3.04e-3) and CD8 effector memory T cells (padj = 0.014), the latter being consistent with increased prevalence of chronic infections (**Fig 3D**). Moreover, Memphians showed a higher frequency and number of pro-inflammatory plasmacytoid dendritic cells (pDCs) (padj = 2.37e-3) (**Fig 3D**).

Substantial variation was also observed between the two cohorts at the RNA level after PBMC stimulation with plain media (an unstimulated control), lipopolysaccharide (a bacterial TLR4 ligand), Pam3CSK4 (a bacterial TLR1/2 ligand), transfected poly(I:C) (representative of RNA virus infection), or transfected poly(dA:dT) (representative of DNA virus infection). For each stimulation condition, samples included 94 Memphians and 94 Nicaraguans. Examples of genes which varied by cohort include, but are not limited to, *IFI16* (a PRR important for herpesvirus detection), *IRF1* and *IRF9* (upstream immune response transcription factors), *TBK1* (a downstream immune signaling protein), *CCL4* and *CXCL2* (chemokines), *CD86* (an antigen presenting cell activation marker and ligand for a co-stimulatory molecule expressed on T cells), *KLRC1* (inhibitory receptor NKG2A, expressed on NK and T cells), and *IFNG* (an antiviral cytokine) (**Fig 4A, Supp Table 3**). These genes represent each step of the immune response and highlight the numerous ways and potential for unique immune profiles to arise in diverse populations. Notably, differential gene expression was stimulation condition dependent, highlighting the need for future studies to investigate immune variation under a variety of immune pressures.

**Figure 4.**
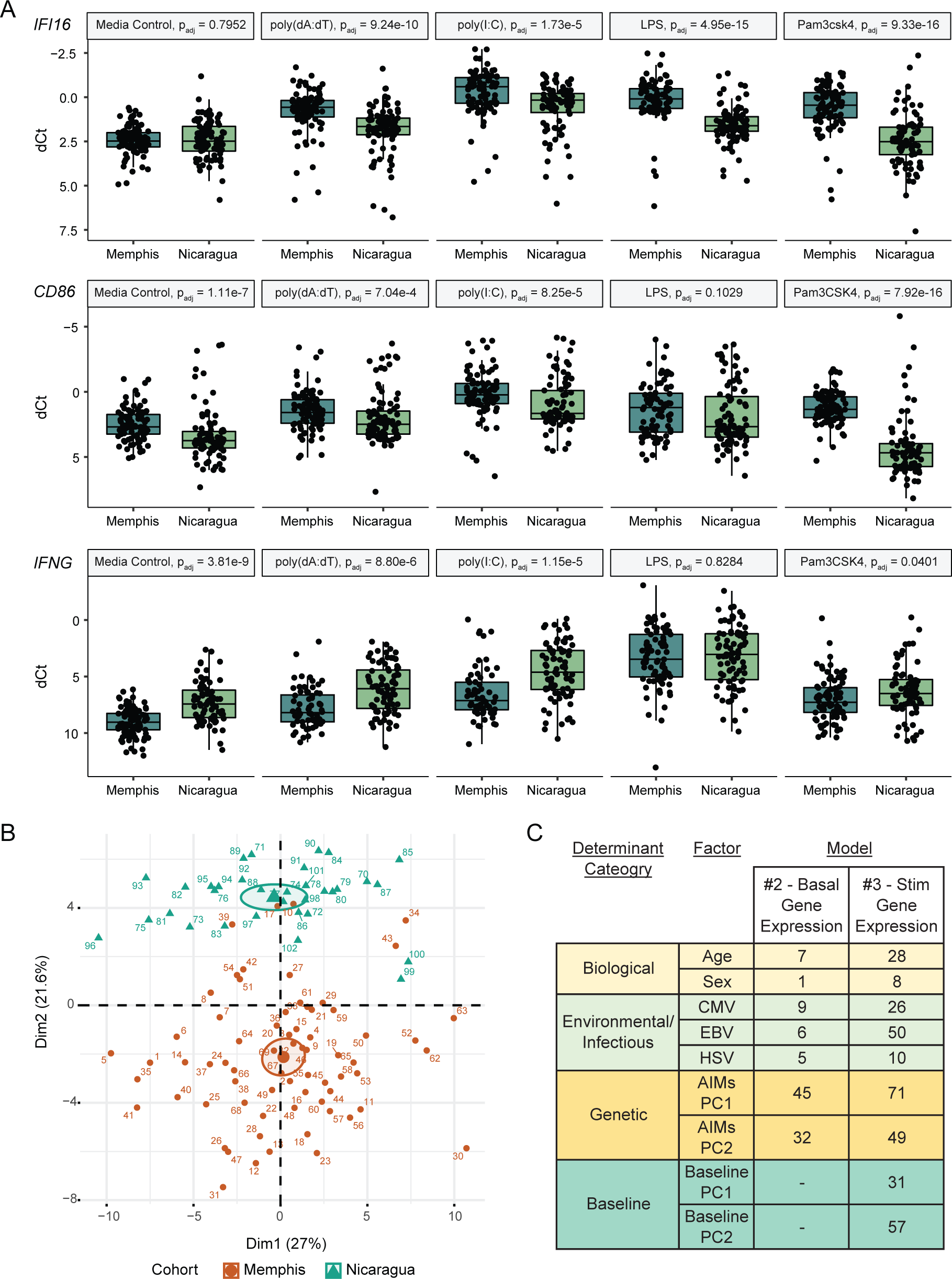
Distinct basal immune profiles mold response to immune challenge. (A) Gene expression by cohort under different stim conditions for *IFI16* (n_AllStim_ = 903), CD86 (n_AllStim_ = 906), and IFNG (n_AllStim_ = 774). Displayed p-values were adjusted for multiple comparisons and are from Mann Whitney U tests. (B) Principal component analysis of baseline (unstim) gene expression colored by Cohort (n = 102). (C) Summary of linear regression Models #2 (n ≈ 188) and #3 (n_AllStim_ ≈ 350) that were utilized to determine factors which contribute to baseline and post stim gene expression variation, respectively (see Summary of Models section for details). Displayed values are the number of genes that a factor significantly affected out of a total 90 genes.

Coinciding with the basal cytokine results (**Fig 3A**), PCA analysis of the media (unstimulated control) gene expression data shows substantial variation in the basal immune transcriptional profile (**Fig 4B**). To determine which factors were driving this variation, linear regression modeling was utilized to account for age, sex, herpesvirus status, and ancestry (Model #2, **Fig 1**). These results show basal immune transcriptional heterogeneity is largely driven by genetic ancestry (affected 62.2% of 90 genes), followed by herpesvirus infection (22.2% of genes), and demographic variables, age and sex (7.8% of genes) (**Fig 4C, Supp Fig 4A, Supp Table 3**). A similar model was utilized to assess determinants of stimulated gene expression, and accounted for stimulant, age, sex, herpesvirus status, ancestry, and the basal immune profile (Model #3, **Fig 1**). Similar to the basal response, ancestry was the most prominent determinant, contributing to statistically significant alterations in 83% of genes examined, the basal transcriptional profile affected 74% of genes (on top of the effects already captured by ancestry), and herpesvirus status affected 66%, while demographic variables (age, sex) affected 31% (**Fig 4C, Supp Fig 4B, Supp Table 3**).

Exploring these transcriptional results further, expression quantitative trait loci (eQTL) were determined to define SNPs that were associated with altered gene expression and demonstrate a means by which genetic effects could contribute to the observed transcriptional heterogeneity. These data were analyzed with linear regression modeling in order to account for: genotypes for immune-related SNPs of interest, genetic based ancestry, age, sex, herpesvirus status, and the stimulation condition (Model #4, **Fig 1**). Of the 94 prioritized candidate SNPs, 36 SNPs in 26 genes (24 cytokines and 2 transcription factors) were significant factors in the expression of their respective genes (**Supp Fig 4C, Supp Table 4**). Notably, 13 of these genes also showed a measurable heritable influence in Brodin et al. at the protein level (GCSF, IFNg, IL10, IL12p70, IL13, IL15, IL6, TNFb, GROa, IL8, MCP1, TGFa, and VEGF).^7^ While further study is required to determine the mechanism behind the eQTLs, these data demonstrate how genetic variation can significantly affect gene expression both at baseline and during immune responses. With respect to immune variation, this would be of most interest when eQTL allele frequencies vary across diverse populations, such as that shown for rs13900 in *MCP1*, which had a minor allele frequency (MAF) of 4% in Memphis and 38% in Nicaragua (**Fig 2D, Supp Table 1**).

### Divergent age-associated influenza cytokine correlates of severity

Results from unstimulated, baseline analyses demonstrate significant immune heterogeneity across diverse populations at the RNA, protein, and cellular level. Further, these data establish that basal immune variation is differentially affected by intrinsic and extrinsic host factors. The cumulative effect of modulation in the basal immune state may lead to unique immune profiles in response to infection. In support of this, the *in vitro* TLR stimulation results show distinct transcriptional responses between the Memphis and Nicaragua cohorts. To further assess the extent to which unique immunophenotypes may occur, we examined how robust immune signatures are across distinct cohorts during acute viral infection *in vivo*.

We determined cytokine correlates of severity in three independent human cohorts of naturally acquired influenza virus infection. Cytokine levels were measured in plasma and/or the nasal wash from influenza positive subjects in the Memphis (n = 165), Baltimore (n = 222), and Taiwan (n = 226) cohorts. Each cohort was independently analyzed and results were compared to identify unique and shared correlates. Utilizing a subset of samples, a previous study in the Memphis cohort found increased levels of nasal wash IFNa2, MCP3, IL6, and VEGF correlated with total, upper respiratory (URT), and/or lower respiratory (LRT) symptom scores. ^33^ Additionally, increased plasma MCP3, IL10, and IL6 predicted hospitalization and were positively associated with symptom scores, and IP10 positively correlated with influenza viral load. ^33^ The current Memphis analysis expands upon this dataset and includes three additional seasons of cytokine data.

There are key similarities and differences amongst the cohorts included in this analysis that are important to consider in interpretation of the results. First, the Memphis cohort is centered around pediatric recruitment, and thus is enriched in children and young adults, though they also enrolled adult contacts. In contrast, the Baltimore and Taiwan cohorts are both adult studies. Additionally, the Baltimore and Memphis cohorts are similar in genetic and geographical background, whereas Taiwan is distinct from the rest. Further, the Baltimore, and Taiwan cohorts each have binary (presence/absence) disease outcome data whereas the Memphis cohort includes both binary and continuous outcomes, the latter of which requires a different statistical approach. Importantly, binary disease outcomes with a high prevalence dilute the specificity of our severity measures, and thus are removed from downstream analyses. Results below are focused on continuous variables or binary variables with a prevalence <40% (range 14-39%, additional details in Study Subjects section of Methods).

All cohorts exhibit increased levels of plasma IL6, TNFa, GCSF, and IL8 in hospitalized, pneumonia, and/or supplemental oxygen patients. This suggests these cytokines are associated with severe influenza disease in general, irrespective of age and geographical location (**Fig 5, Supp Fig 5, Supp Table 5**). However, these analytes may reflect severe (respiratory) viral infection more broadly, as they have also been associated with severe COVID-19.^39–42^ Cytokine profiles as a whole, in both the plasma and nasal wash, are distinct by age groups, possibly reflecting unique pathophysiologies. The severe pediatric response is characterized by global hyper-inflammatory cytokine levels, whereas the severe adult response shows largely decreased cytokine levels with a selective inflammatory response of the aforementioned analytes (**Fig 5, Supp Fig 5, Supp Table 5)**. Notably, although the directionality is similar among significant results in the adult cohorts, only decreased IFNg was shared, and thus represents the most robust correlate for this age group. All additional plasma cytokine correlates of severity in these adult populations were also associated with lower cytokine levels, albeit specific to one of the adult cohorts (**Fig 5, Supp Fig 5, Supp Table 5**).

**Figure 5.**
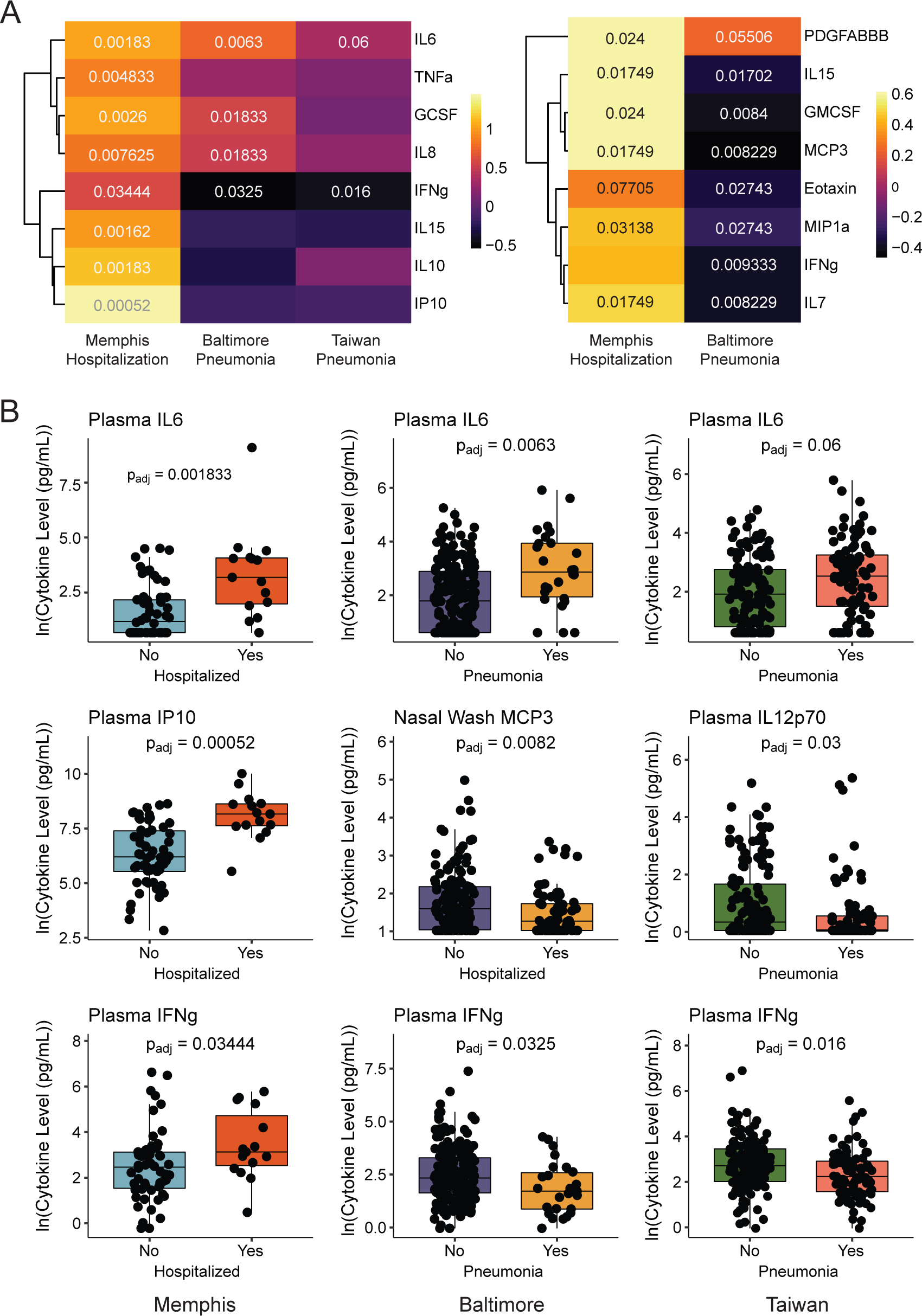
Acute cytokine profiles delineate two age-associated signatures of influenza disease severity. (A) Heatmap of highlighted results from plasma (left) and nasal wash (right) cytokine correlates of severity analysis, with significant adjusted p-values displayed, for the Memphis (n = 165), Baltimore (n = 222), and Taiwan (n = 226) cohorts. (B) Boxplots show example associations between plasma or nasal wash cytokine levels and symptom severity during influenza infection for each cohort. Presented p-values are from Mann Whitney U tests, and results were adjusted for multiple comparisons by controlling the FDR (< 0.1).

These data demonstrate correlates of severity can differ across distinct populations and highlight the need for future work to simultaneously examine immune signatures of health and disease in diverse populations for a more comprehensive analysis and identification of robust therapeutic targets. Additionally, studies which aim to identify cytokine correlates should utilize a broad panel, as focusing on a few select analytes is insufficient to characterize the overall cytokine profile. These results also suggest that a “one size fits all” therapeutic approach may not be efficacious, as distinct immune profiles may exist by extremes of age and likely require different treatments. Multiple severity associated immunotypes were also recently observed in a study of SARS-CoV-2 cellular responses, wherein 3 distinct cellular immunophenotypes were identified for severe COVID-19.^43^

### Age specific herpesvirus associations with influenza severity

In addition to immune correlates of disease severity, independent studies have identified associations between age, sex, and CMV and the outcome of vaccination or infection with influenza in humans.^33,44,45^ To expand upon these findings, we examined disease outcome utilizing a linear regression model which accounts for age, sex, CMV, EBV, and HSV serostatus, as very few individuals are only infected with CMV (**Fig 2E-F, Supp Fig 1**).

In Memphis, the pediatric cohort, CMV serostatus was the only significant variable and it was associated with improved outcome. Patients with a positive serostatus are enriched in the non-severe subjects, and show a decrease of 29.1 ± 10.93 (p = 0.0103) in peak lower respiratory symptom severity score (range = 0 to 176) (**Fig 6A, Supp Table 6**). Conversely, in the adult Baltimore cohort, a positive CMV serostatus was associated with increased risk of pneumonia (OR = 2.904, p = 0.0457) or hospitalization (OR = 4.966, p = 5.72e-5) (**Fig 6A, Supp Table 6**).

**Figure 6.**
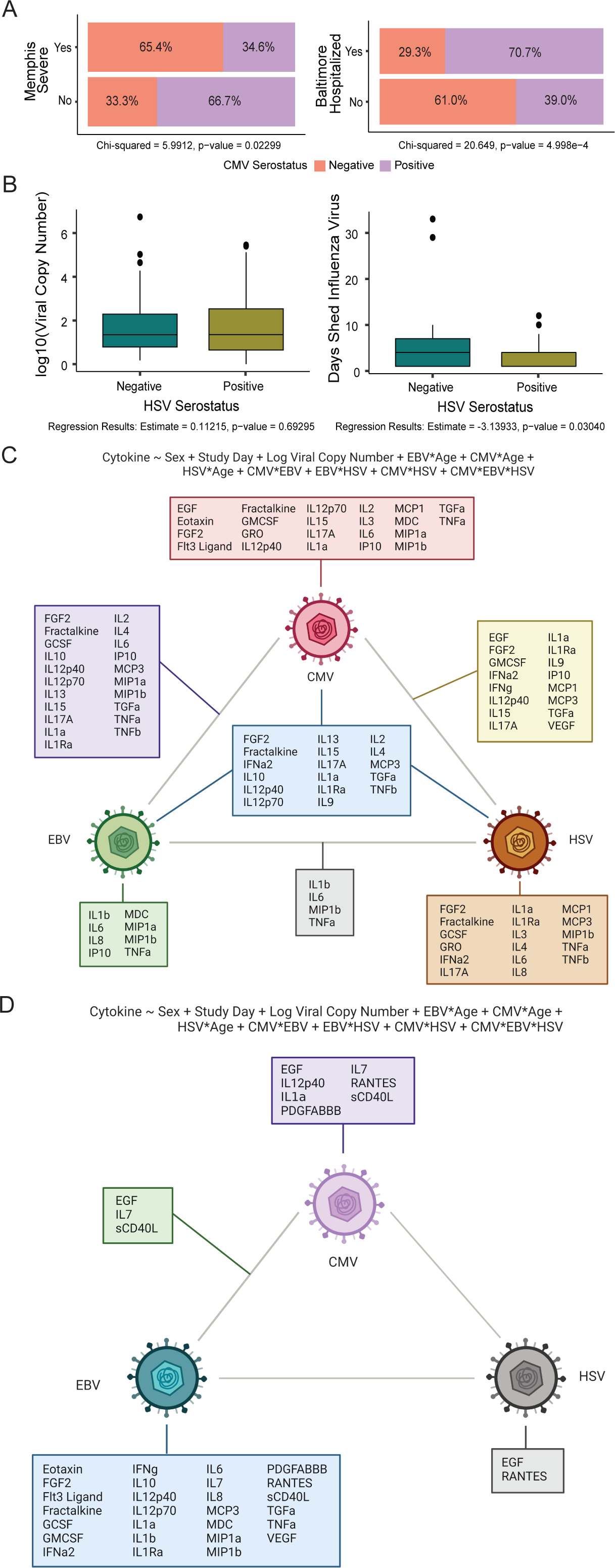
Herpesviruses have unique and age specific effects on influenza disease severity. (A-B) Significant herpesvirus associations with influenza disease severity. Severe Memphis was categorized based on lower respiratory symptom severity scores. (C-D) Schematic of herpesvirus effects on acute cytokine levels in plasma (C, n ≈ 200) and nasal wash (D, n ≈ 300) based on linear regression modeling results (models displayed above diagrams). Cytokines listed have an adjusted R^2^ value > 0.08 and FDR < 0.05.

Aside from symptom severity, the Memphis cohort included two additional measures of the quality of the immune response to influenza infection - influenza viral load was determined using qRT-PCR (quantitative reverse transcription polymerase chain reaction) and functional antibody titers were measured using microneutralization assays. To analyze these data, linear regression modeling was utilized to assess the effect of biological (age, sex) and infectious (herpesvirus serostatus) factors, while accounting for study day (when applicable). Neither CMV, EBV, nor HSV had any effect on influenza viral load (p = 0.66, 0.39, and 0.69, respectively) (**Fig 6B, Supp Table 6**). However, a positive HSV serostatus is associated with a decrease in duration of influenza virus shedding by approximately 3.14 ± 1.42 days (p = 0.0304, **Fig 6B, Supp Table 6**). These results suggest that HSV co-infection may alter viral clearance mechanisms, such that it decreases the days an individual sheds influenza virus. Although, there was no detected effect on influenza viral load, one challenge with observational human studies is unknown true day of infection and the viral load upon initial infection.

Analysis of microneutralization titers show that a positive CMV serostatus is associated with an increase in functional antibody titers by approximately 144 ± 54.50 (p = 0.0086) (**Supp Table 6**). Additionally, male sex is associated with an increase in antibody titer of 199 ± 60.70 (p = 0.00112) and titer decreases by 105.04 ± 30.60 per standard deviation unit change in age (p = 0.00067). These results are consistent with previous observations by Furman et al. which show that CMV seropositive young adults exhibit augmented antibody responses to Fluzone vaccination compared to CMV seronegative.^44^

Collectively, these data suggest a dichotomous, age-specific role of CMV on disease severity. This is consistent with prior work which has shown CMV improves vaccine responses in young populations, but increases frailty in the elderly.^44,46–48^ Initial benefits may be mediated by trained innate immunity, in which constant surveillance of chronic herpesvirus infection (in order to mitigate reactivation) necessitates altered epigenetic and transcriptional profiles which confer augmented response to restimulation.^49^ However, the cumulative effects of CMV on immune aging over time, such as through inflammaging and memory inflation, may eventually exceed beneficial effects and lead to impaired, immunosenescent responses and, subsequently, poorer outcome.^50–52^ Future study is required to delineate the extent to which trained innate immunity, CMV associated immunosenescence, or an alternative mechanism is the cause of these differential influences on heterologous immunity.

### Differential contributions of host genetics and herpesviruses to acute immune variation

The correlates of severity analysis demonstrates immune heterogeneity in acute immune responses, and thus variation in putative therapeutic targets. Moreover, it identified biological and infectious associations with influenza disease outcome. Having observed biological and infectious contributions to basal cytokine levels, we sought to determine whether the association between disease severity and these factors may be due, at least in part, to their effects on cytokines, particularly those that were identified as a correlate of severity. These results will also provide insight into whether such analyses would be useful in the identification of optimum therapeutic targets by determining the extent to which there are differential contributions of intrinsic and extrinsic host factors on cytokines of interest.

To achieve this, plasma and nasal wash samples were used from 83 subjects with naturally acquired influenza infection in the Memphis cohort from time points throughout the duration of infection. A total of 38 and 41 cytokines were measured in plasma (n = 237) and nasal wash (n = 325), respectively. Since eQTLs were identified for a subset of analytes, this analysis begins with an assessment of the effects of age, sex, and herpesviruses on cytokine levels, while accounting for influenza infection related variables that can contribute to the immune state (Models #5-6, **Fig 1**). This is followed by a second regression analysis, focused on cytokines with an eQTL identified in the baseline analyses, which expands upon the previous model and incorporates genetics by adding AIMs PCA values and SNP genotypes (Models #7-8, **Fig 1**).

For all analytes, a linear regression model was used to take into account the effects of: age, sex, study day, influenza viral load, each herpesvirus alone, any interaction between a herpesvirus and age, and any interaction between herpesviruses (Models #5-6, **Fig 1**). These data show each herpesvirus has its own unique signature of immune modulation that is location specific. In plasma (**Fig 6C, Supp Fig 6**), CMV alters the most cytokine levels independently, which coincides with its primary site of latency in cells of the myeloid lineage. Furthermore, herpesvirus interactions are most prevalent in the plasma, versus in the nasal wash, where independent herpesvirus effects are more common. In the nasal wash (**Fig 6F, Supp Fig 7**), the herpesvirus with the most independent effects is EBV, which predominantly infects the nasopharynx region. In the plasma, the adjusted R^2^ value reached 0.358 (or 35.8%) for IL10, and herpesvirus interactions are generally associated with the strongest effects (i.e. magnitudes) on cytokine levels as compared to the effects of independent herpesviruses or other variables (i.e. age, influenza viral load) (**Supp Fig 6)**. In the nasal wash, the adjusted R^2^ value reached 0.332 (or 33.2%) for RANTES, and three factors accounted for most effects on cytokine levels - viral load, age, and EBV, with the strongest effects from the latter two (**Supp Fig 7**).

**Figure 7.**
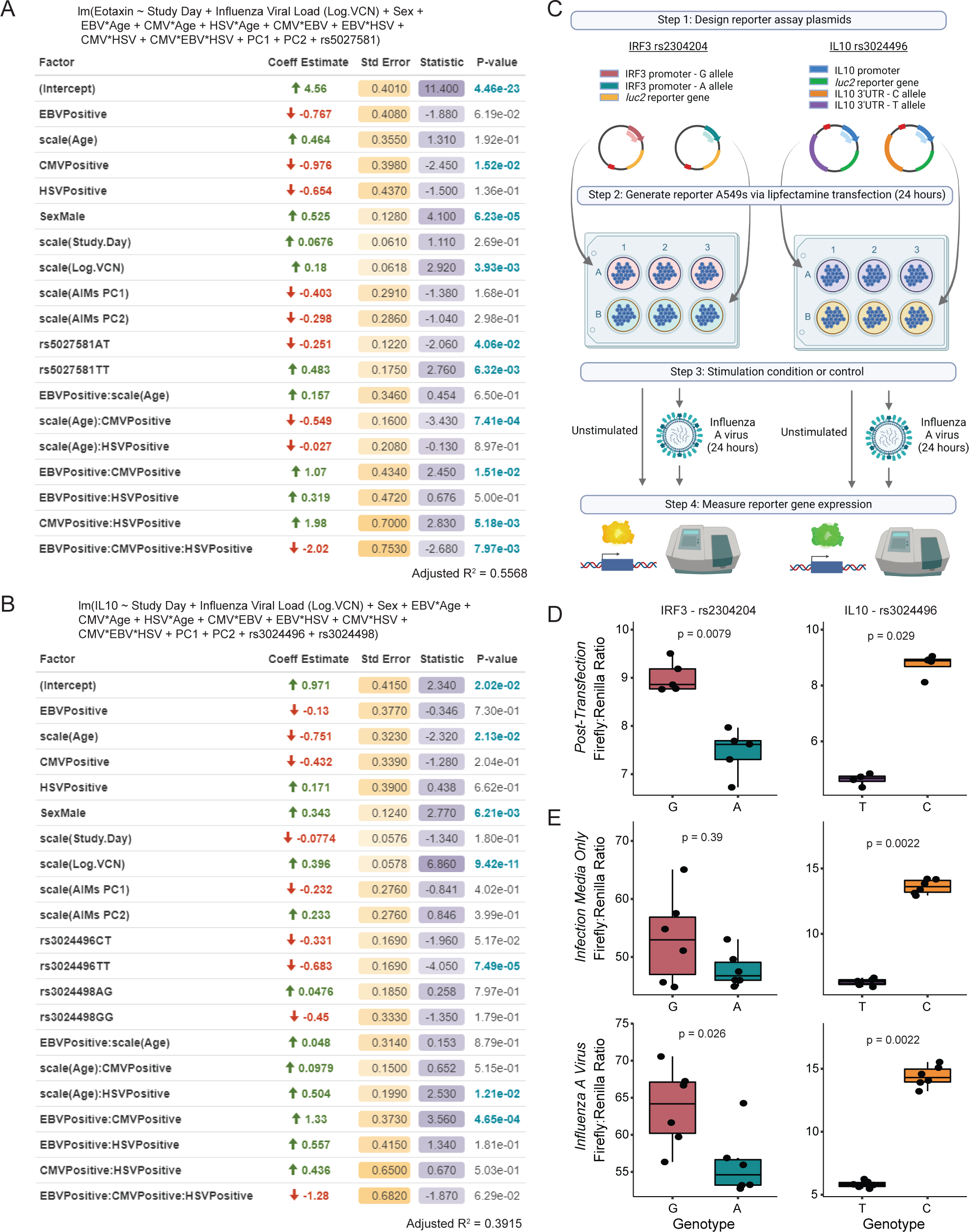
Transcription and protein level of IL10 is modulated by rs3024496. (A-B) For cytokines with an eQTL, a linear regression model was performed including: sex, age, herpesvirus (HV) serostatus, interactions between HV serostatus and age, interactions between HVs, study day, viral load, AIMs PC1 and PC2, and eQTL SNP genotype (n ≈ 200). The top two results, based on adjusted R² value, are shown and include the coefficient estimate of the change in the dependent variable per unit change in the independent variable (Coeff Estimate), the corresponding standard error (Std Error), t score (Statistic), and p-value (highlighted when significant). (C) Schematic of luciferase assays. Boxplots compare Firefly:Renilla ratios from unstimulated (D) or influenza virus infected (E) luciferase assays for A and G alleles of IRF3 rs2304204 (left), and T and C alleles of IL10 rs3024496 (right). Presented p-values are from Mann-Whitney U tests.

For the 23 cytokines which had eQTLs and protein level data, a second regression model was performed for plasma and nasal wash, which included each factor previously described (age, sex, herpesviruses, study day, viral load), in addition to the AIMs principal components 1 and 2 (genetic based ancestry) and the genotype for each SNP that significantly correlated with the respective cytokine’s gene expression (Models #7-8, **Fig 1**). GCSF and its SNPs which trended toward a correlation were also included because fewer samples had detectable gene expression (**Supp Table 7**). In the plasma, when there was an effect from incorporating genetics into the cytokine regression models, it generally increased the amount of variability accounted for by the model; however, the nasal wash models generally decreased. This is likely due, at least in part, to the fact that eQTL status was determined from stimulated PBMCs as opposed to nasal wash cells, which are at the site of infection, have a distinct milieu, and are exposed to the external environment. Eight out of 24 cytokine models had a significant genetic component and increased variability accounted for, two of which solely included a significant AIMs PC without a significant association with a curated SNP (**Supp Table 7**). These cytokines include chemokines (Eotaxin, MDC, MIP1a), immunomodulatory IL10, NK and memory T cell proliferation inducing IL15, and growth factors (FGF2, GCSF, TGFa). The biggest change in adjusted R^2^, upon inclusion of genetic factors, occurred in Eotaxin with a 43.35% increase in the variability accounted for by the model, for a final adjusted R^2^ value of 0.5568 (**Fig 7A**). Significant factors in the Eotaxin model include: sex, influenza viral load, SNP rs5027581, CMV and its age interaction, the CMV and HSV interaction, the EBV and CMV interaction, and the interaction of EBV, CMV, and HSV. The next highest adjusted R^2^ was 0.3915 (39.15%) for plasma IL10 and significant factors include age, sex, viral load, SNP rs3024496, the HSV and age interaction, and the EBV and CMV interaction (**Fig 7B**).

### Validation of the model: rs3024496 genotype modulates IL10 expression

To follow up on genetic associations from our analyses, we selected two associated SNPs with potentially functional consequences for *in vitro* assays to assess effects on promoter activity. SNP rs3024496 in *IL10* was chosen because IL10 is a correlate of severe influenza infection, and its genotype prevalence varied across diverse populations (Memphis: 39% TT, 47% CT, 14% CC; Nicaragua: 67% TT, 32% CT, 1% CC). Of note, two SNPs within the gene encoding IL10 were identified as eQTLs (rs3024496, rs3024498). However, including both SNPs in the IL10 regression model served as a conditional SNP analysis and suggested there was only one independent signal resulting from rs3024496. SNP rs2304204 in *IRF3* was also selected because it is a transcription factor upstream of antiviral responses, with varying allele prevalences across cohorts (Memphis: 40% GG, 40% AG, 20% AA; Nicaragua: 11% GG, 45% AG, 44% AA). Additionally, an independent analysis using the GTEx Portal confirmed our transcriptional results and showed rs2304204 is an eQTL in tissues such as the thyroid (p = 1.1e-19) and lung (p = 0.000027).

For each target, two luciferase reporter assay plasmids were created, which only differed in sequence at the SNP site (**Fig 7C**, detailed in Methods). Unstimulated A549s exhibited higher luciferase expression when the C allele was present for IL10-rs3024496 and when the G allele was present for IRF3-rs2304204 (**Fig 7D**). Results were consistent 24 hours post infection with influenza A virus and showed a higher magnitude of differential activity (**Fig 7E**). Coupled with the eQTL findings, these results confirm rs3024496 is a functional SNP directly affecting IL10 promoter activity and coincide with the association predicted by the regression model - the CC genotype was associated with higher plasma IL10 than the CT and TT genotypes which had coefficient estimates of −0.3315 and −0.6830, respectively, when compared to the reference CC genotype (**Fig 7B**). SNP rs2304204 in *IRF3* was also verified as a functional SNP with coinciding directionality. This finding, coupled with the varying allele prevalence across diverse populations, demonstrates a mechanism by which broad transcriptional changes could occur, both at baseline and in response to immune challenge, as was observed in this study (**Fig 4**). Together, these results demonstrate the integrative manner in which biological, infectious, and genetic factors can influence variation in cytokines at the protein level during naturally acquired influenza infection. The observation that these effects occur in proteins identified as correlates of severity, highlights how variation in these factors can lead to distinct immunophenotypes during acute infection and the potential impact on infectious disease outcome. Collectively, this underscores the need for future studies to search for correlates of protection or severity across diverse populations and to consider the factors which contribute to their variation in order to identify the number of severe immune profiles, the most broadly applicable treatment, and practical therapeutic options (i.e. those more affected by factors we can control or target).

## Discussion

Studies of human immune variation thus far suggest genetics and age contribute to 20-40% and 5% of immune variance, respectively, leaving 55-75% possibly due to sex or environmental influences.^53^ However, studies to date often focus on a single ancestral group, geographical location, immunological measure, and/or age range, and are sometimes limited in inclusion of both sexes.^4,6–8,20,21,54–56^ Often, this is in an attempt to limit additional variance attributed to determinant categories that are not the focus of a given study. While statistically beneficial and still informative, it is impossible to ascertain the scope of immune variation when studies are focused on specific subsets of the human population and this limits the applicability of the conclusions that are drawn. Moreover, this can obscure phenotypes, leading to false positive or negative associations, and inconsistent results across studies that ultimately present a barrier to translating results from the laboratory to effective, broadly applicable therapeutic treatments. Lastly, comparisons of relative importance can be better understood with simultaneous assessment of multiple determinant categories, and with increasing technologies for multiparameter assessment, this approach is becoming more tractable, as demonstrated by this study.

The demographics, immune measures, and analytical approach make this study unique from prior work. First, the populations included are diverse in ancestral background, geographical location, age, and sex. Second, immune measures were determined at baseline and during immune challenge (both *in vitro* and *in vivo*), and include cell phenotypes, cytokines (both circulatory and at the site of infection), and gene expression in response to PRR ligands representative of both bacterial and viral infection. Third, the analytical approach utilized linear regression modeling which allowed us to simultaneously account for and quantify the unique and interactive effects of biological, infectious, and genetic factors. Fourth, our genetic approach was based on ancestry informative markers and a targeted list of candidate immune-related SNPs. This allowed for increased power compared to GWAS studies by limiting the impact of multiple comparisons and provided clearly defined genetic factors that can be further explored in mechanistic studies that aim to identify causal factors compared to heritability scores which are unable to provide direct functional insight.

Investigation of baseline immunity shows differences in cellular phenotypes, cytokine levels, and the basal transcriptional profile. Heterogeneity in baseline gene expression was primarily influenced by genetic ancestry, followed by (in order of importance) herpesvirus infection, age, and sex. Activation and acquisition of effector function is dependent on both the microenvironment and differentiation state of leukocytes, and variation in these facets of basal immunity were reflected in response to pathogen-like challenge. Stimulation with bacterial or viral ligands showed differential transcriptional responses across the Nicaragua and Memphis cohorts in genes important for each aspect of acute immunity. Differences between cohorts were often dependent on the stimulatory conditions, suggesting that disparate immunophenotypes can be context/infection dependent and further investigation of immune variation under a wide spectrum of immune pressures is needed. Furthermore, the transcriptional response to stimulation is largely affected by genetic ancestry. Remaining determinants, in order of impact, include the basal transcriptional profile, herpesvirus status, and demographic variables, age and sex. Together, these results highlight the substantial impact of genetics on immunity, underscore the need for more studies of diverse ancestral backgrounds, and provide a greater understanding of the relationship between the baseline immune state and the magnitude and quality of acute immunity.

In further support of unique immune signatures across distinct populations, analysis of plasma cytokines in three independent human cohorts of influenza infection showed two influenza disease severity profiles, largely driven by age, wherein the severe pediatric response exhibits global hyper-inflammatory cytokine levels and the severe adult response exhibits generally decreased cytokine levels, with increases of select cytokines (IL6, TNFa, GCSF, IL8). This indicates different therapeutic approaches may be needed and underscores the need for future work to search for therapeutic targets across diverse populations in order to identify the most robust and broadly applicable options. Such studies would advance the scientific and medical communities closer to achieving personalized medicine by providing valuable insight into the number of distinct severity profiles, the appropriate method to treat severe populations/profiles, and the minimum number of therapeutic approaches required to effectively treat patients of all demographics.

Results from individual cytokine regression models, both at baseline and during infection, show each category of independent variables contributes to immune variation, these effects are not mutually exclusive, the magnitudes of modulation vary by the cytokine of interest, and there are interactive effects between and within categories. For example, herpesviruses have unique and interactive effects on cytokine levels, suggesting independent and collective mechanisms of immune modulation and emphasizing the need for future studies to include additional herpesviruses in their experimental design. Interestingly, the effects of herpesviruses at baseline are much smaller compared to the effects during acute infection with influenza. This may be due, at least in part, to the limited transcriptional profile of herpesviruses during latency, whereas increased transcriptional activity, perhaps in response to immune cell differentiation or signaling, would allow for greater interference due to increased viral protein production. Indeed, established methods for reactivating herpesviruses include stimulation with PRR ligands and differentiation of myeloid cells.^57,58^

Of the 23 cytokine models with acute infection (study day, viral load), genetic (SNP genotypes, AIMs), chronic infectious exposure (herpesvirus serostatus) and biological (age, sex) factors, 8 had a significant genetic component and improved model performance, each of which were identified as a correlate of influenza disease severity. Functionally relevant SNPs, identified in eQTL and protein cytokine level regression analyses, included SNPs with different allele frequencies across populations and SNPs with similar allele frequencies across populations but variations within populations, highlighting genetic contributions to both inter- and intra-population immune variation, respectively. Importantly, variation in genotype prevalence across or within populations may lead to different immune responses and subsequent immune outcomes if the altered protein is a correlate of severity. For example, SNP rs3024496 was identified as an eQTL for IL10, its distribution of genotypes varied between Memphis and Nicaragua, and prior work in infected subjects from the Memphis cohort found IL10 to be a correlate of hospitalization and symptom severity during influenza infection.^33^ Moreover, IL10 had the second highest adjusted R^2^ value in the infected regression model, and factors which contributed to its variation, at the protein level, included SNP rs3024496, age, sex, viral load, and herpesviruses. Luciferase assays further support the modeling results and show the functional impact of rs3204496, such that the C allele is associated with higher promoter activity when unstimulated or infected with influenza A virus, with the latter condition indicating a higher magnitude of differential gene expression. This demonstrates the powerful implication of genetics on the overall immunophenotype across ancestrally distinct populations, and supports recent work which highlights the value in and need for increased studies of diverse, particularly multiethnic, populations to improve our understanding of human immunology.^25,59,60^

It’s postulated that immune variation can be modeled as an “immunological landscape”, with individuals lying at different coordinates according to their immune profile, this provides a framework for the interaction of genetic, environmental, and biological factors in shaping the human immune response.^53^ In the initial description, it was suggested that the starting position of an individual within the landscape is largely due to genetics, whereas age changes this position in a consistent and predictable manner across subjects, and local environment may drive convergence between two individuals if shared (such as cohabitation) or diverge two individuals if they are from distinct environmental backgrounds.^53^ The results presented here suggest an even more complex and multidimensional immune space with dynamic landscapes depending on the immune pressure and interactions between variables which change the position of an individual, in addition to single variable influences. Our results show there are clusters of distinct immunophenotypes across this landscape, both at baseline and during infection. Importantly, the observation that gene expression differences between two diverse populations depended on the stimulatory condition suggests that the immunological landscape is context dependent, thus unique immunophenotype profiles and clusters may exist during infection that were not observed at baseline. Additionally, unique correlates of severity across diverse populations and shared correlates in cohorts with common features (e.g. age range) suggest distinct clusters of immune profiles are associated with differential illness outcomes and more than one severity-immunotype cluster may exist for a given disease. These observations, and others presented here, likely would have been obscured if the study solely focused on a single population, especially if biased in a given demographic. This underscores the need for future studies to include diverse subjects when examining immune variation, whether at baseline or during infection. While it is challenging to conduct studies that are diverse in all desired categories, this study shows we can leverage multiple cohorts enriched in specific target populations and conduct comparative analyses to gain insight into overlaps or unique signatures of immunity as it relates to immune competence and disease severity.

### Limitations of the Study

Although simultaneous investigation of multiple cohorts provided novel insight, there are several limitations in this study. The independent designs of each cohort and differences in study endpoints affected sampling and data collection. One of the baseline cohorts collected serum, as opposed to plasma; however, we believe the impact was minimal, as it only applies to basal analyses, and a study which compared plasma versus serum cytokine levels, using Luminex multiplex assays, shows results are well correlated.^61^ Differences in sampling types, timing, and quantities limited baseline and acute infection PBMC availability for several cohorts. Thus, we were unable to include acute cell phenotypes and connect baseline immune profiles with acute immunophenotypes. Additionally, multiple approaches were utilized for symptom reporting and clinical data collection, which required different statistical approaches for correlates of severity analyses and limited our ability to include other factors, such as body mass index. Lastly, there may be unknown differences in handling and/or storage across the multiple study sites, which we attempted to account for by including “Cohort” as a factor in our models, when appropriate.

## Supplemental Item Legends

**Supplemental Figure 1.**
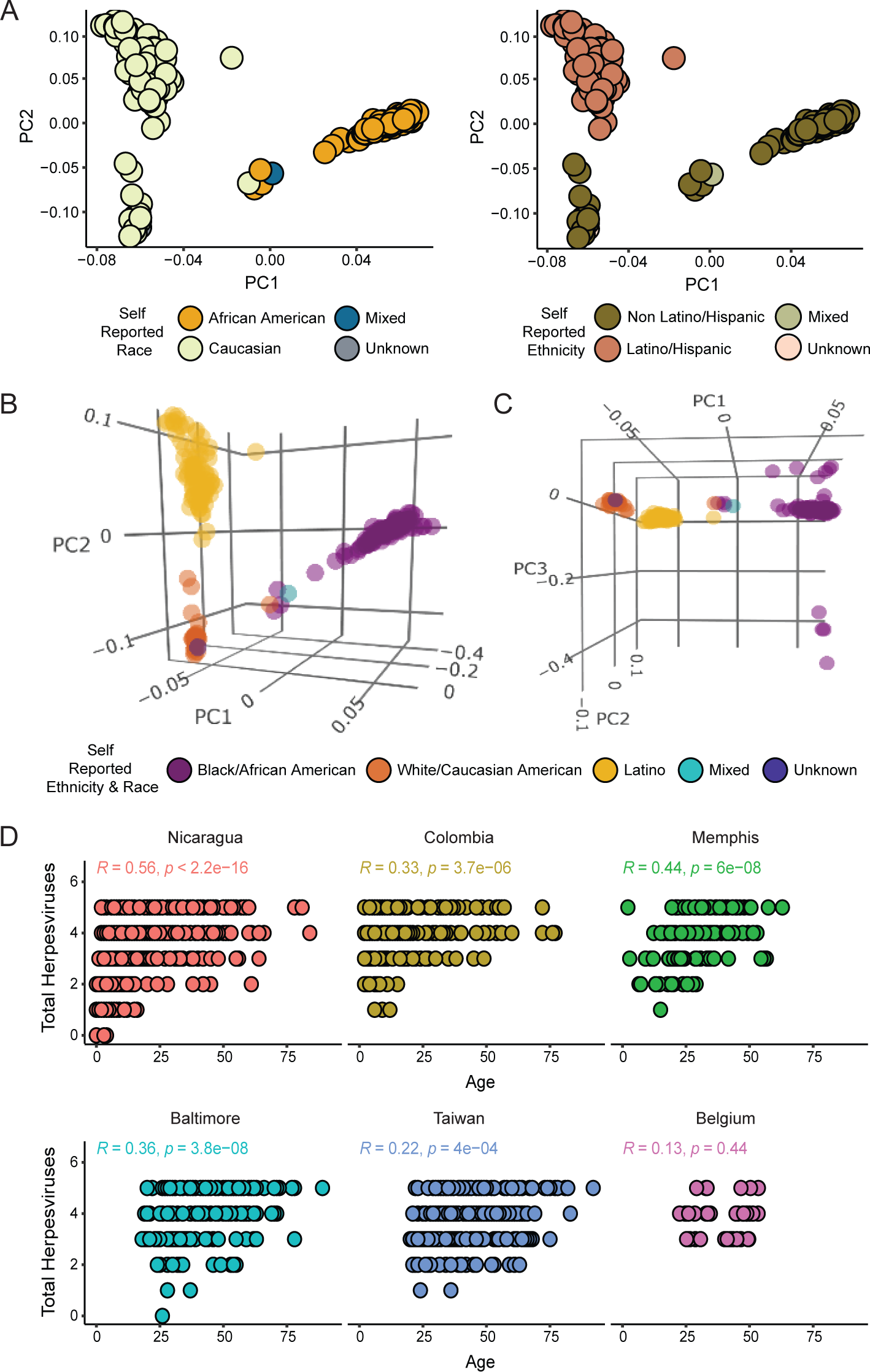
Explanatory variables by Race/Ethnicity and age. (A-C) Principal component analysis of AIMs colored by Race and/or Ethnicity. (D) Total number of herpesviruses by age, separated by cohort.

**Supplemental Figure 2.**
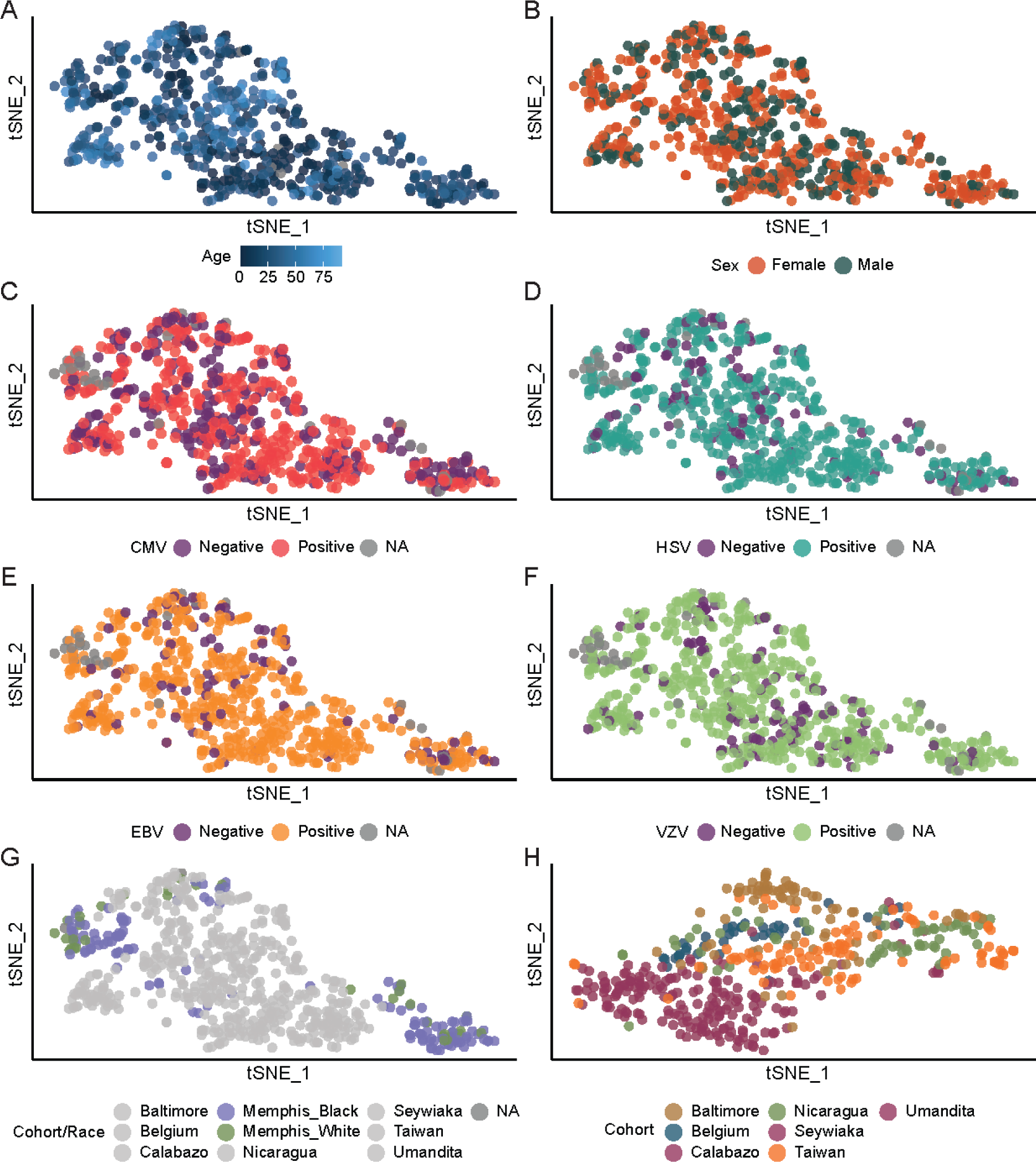
Cohort tSNE clusters are not associated with demographics or chronic virus infection. Basal cytokine tSNE plot colored by age (A), sex (B), herpesvirus serostatus (C-F), and Race (G, for Memphis only, all other cohorts in gray). (H) Basal cytokine tSNE with Memphis cohort data removed.

**Supplemental Figure 3.**
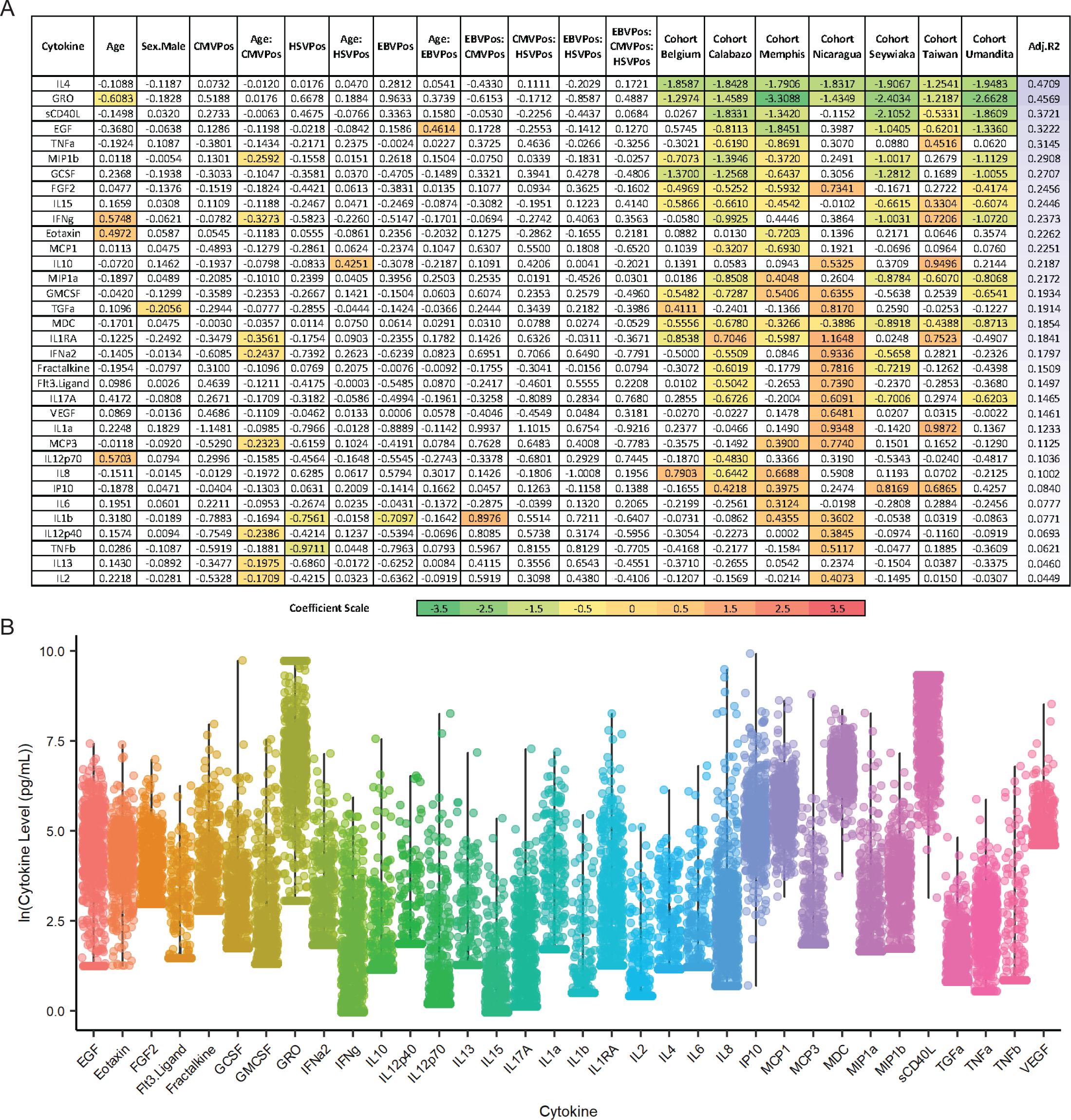
Heatmap summary of baseline cytokine regression models. (A) Summary of coefficient estimates for each factor included in the baseline plasma/serum linear regression model. Cytokine models are listed in decreasing order of the adjusted R^2^ value (shown on the far right) and coefficients are colored if FDR < 0.05. Red is associated with an increase in the cytokine level per unit change in the factor, and green is associated with a decrease in the cytokine level. (B) Basal plasma/serum cytokine level distributions.

**Supplemental Figure 4.**
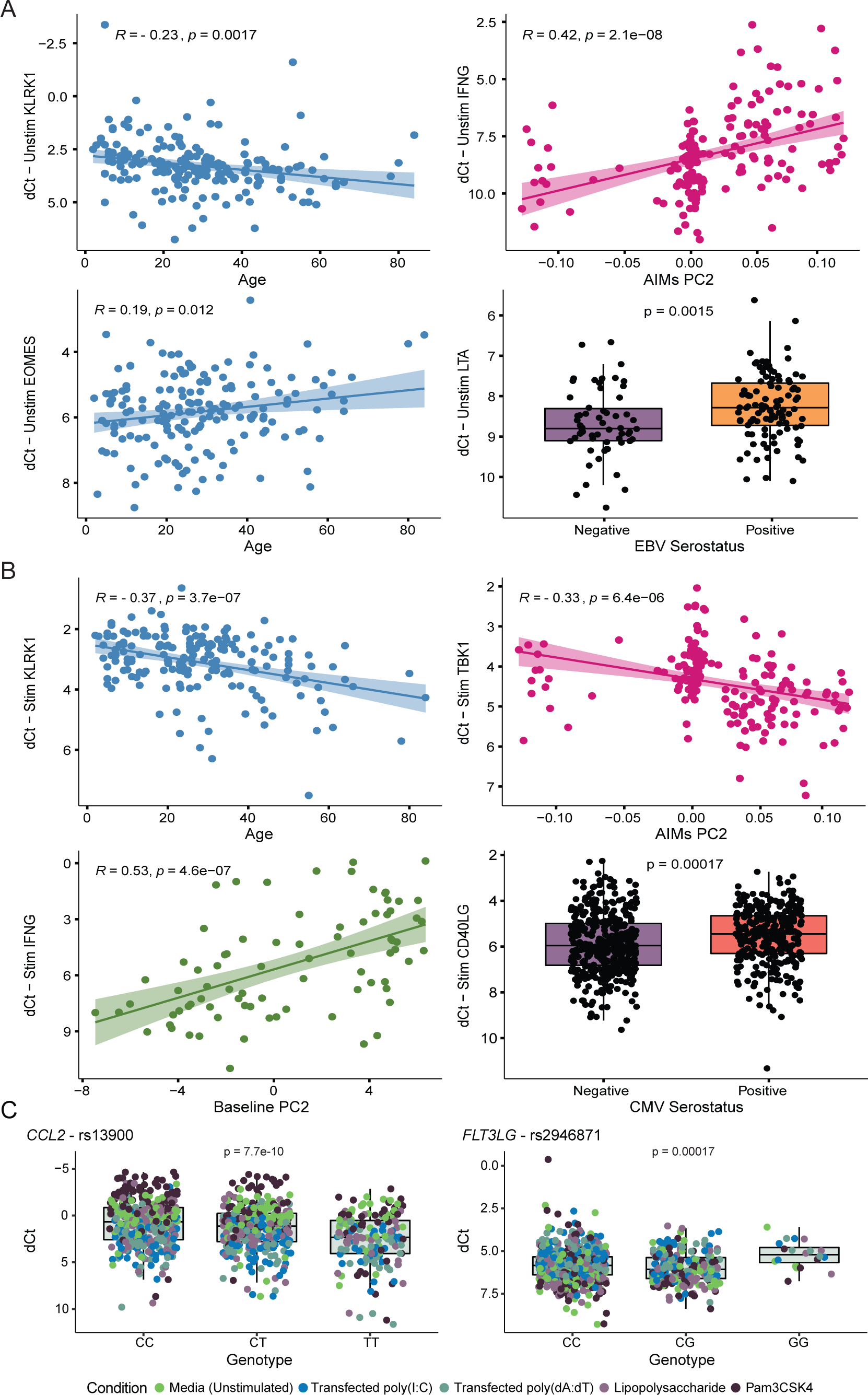
Highlights of gene expression associations with independent variables. (A-B) Example of associations based on results from Model #2 (A, determinants of baseline gene expression) and Model #3 (B, determinants of stimulated gene expression). The y-axis is gene expression and the x-axis is the independent variable (age in years, herpesvirus serostatus, ancestry informative marker principal component, baseline gene expression principal component). (C) Examples of eQTLs. *MCP1* (n = 833) and *FLT3LG* (n = 868) gene expression by genotype for rs13900 and rs2946871, respectively.

**Supplemental Figure 5.**
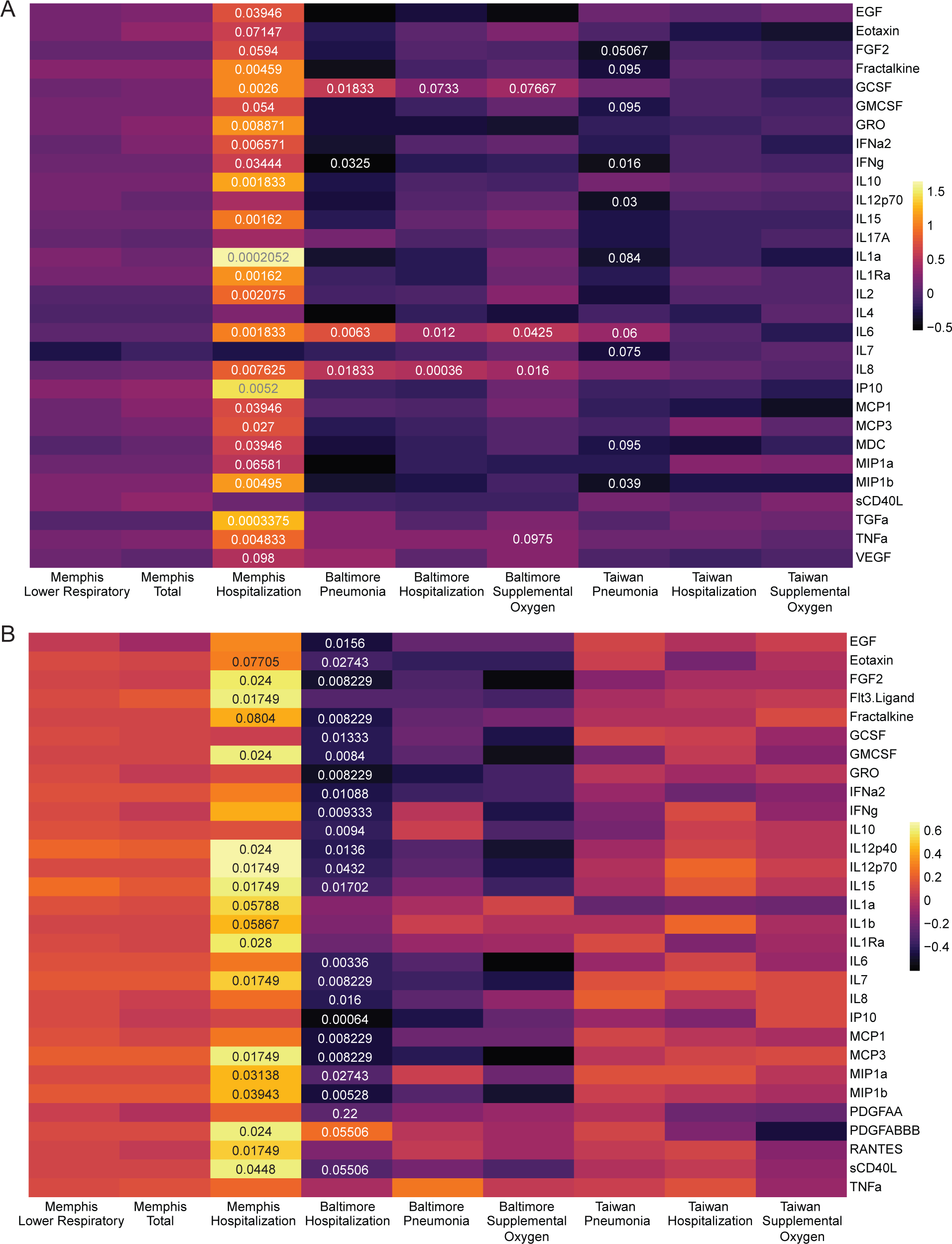
Plasma and nasal wash cytokine correlates of severity. Heatmap of plasma (A) and nasal wash (B) cytokine correlates of severity analysis, with significant adjusted p-values displayed. Colors are based on correlation coefficients or a Cohen’s D effect size, depending on whether a symptom is continuous or binary, respectively. Presented p-values are from correlations (Spearman and Pearson) or Mann Whitney U tests, and results were adjusted for multiple comparisons by controlling the FDR (< 0.1).

**Supplemental Figure 6.**
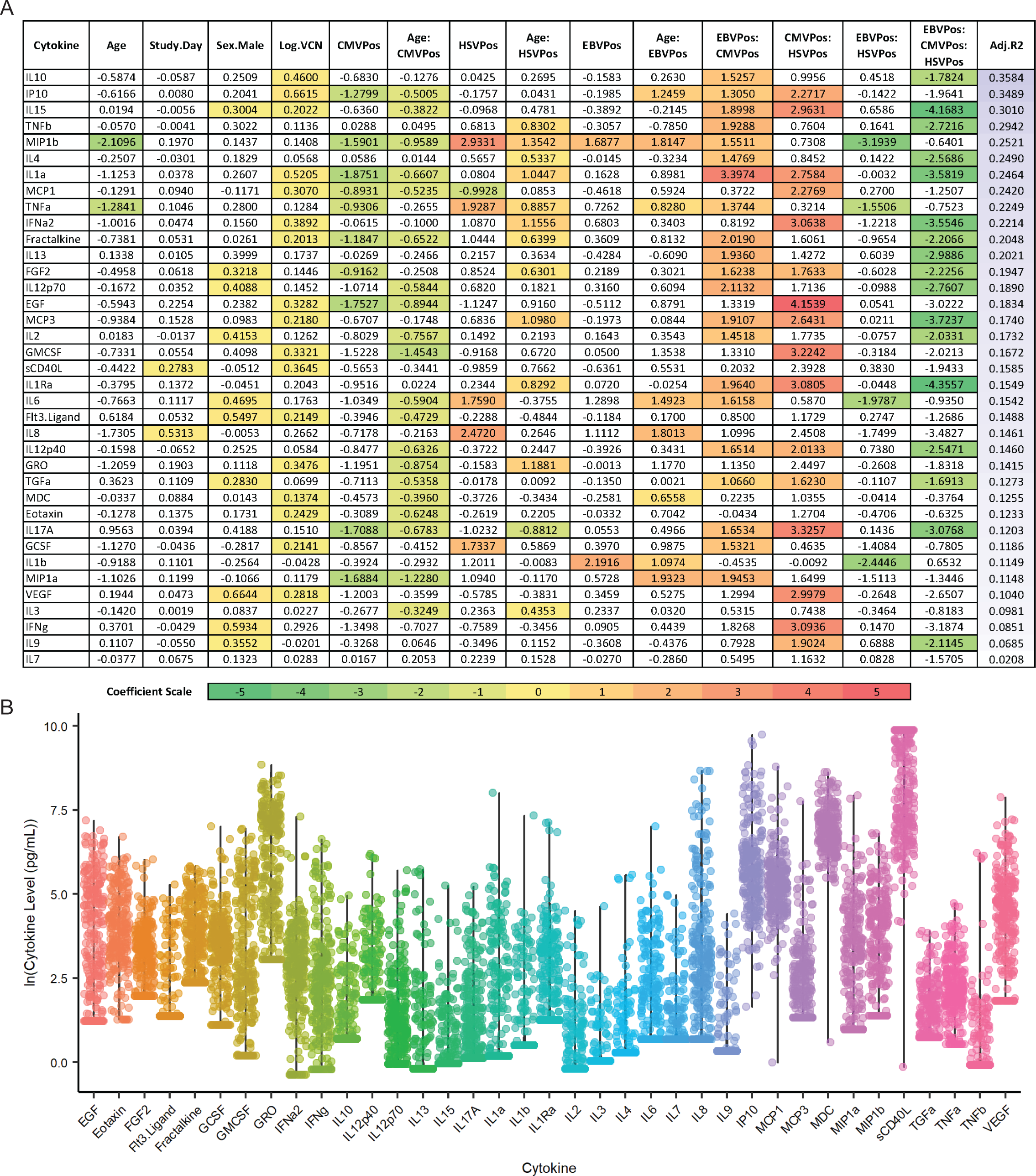
Heatmap summary of acute plasma cytokine regression models. Samples were from influenza infected subjects. (A) Summary of coefficient estimates for each factor included in the infected plasma linear regression model. Cytokine models are listed in decreasing order of the adjusted R^2^ value (shown on the far right) and coefficients are colored if FDR < 0.05. Red is associated with an increase in the cytokine level per unit change in the factor, and green is associated with a decrease in the cytokine level. (B) Acute plasma cytokine level distributions.

**Supplemental Figure 7.**
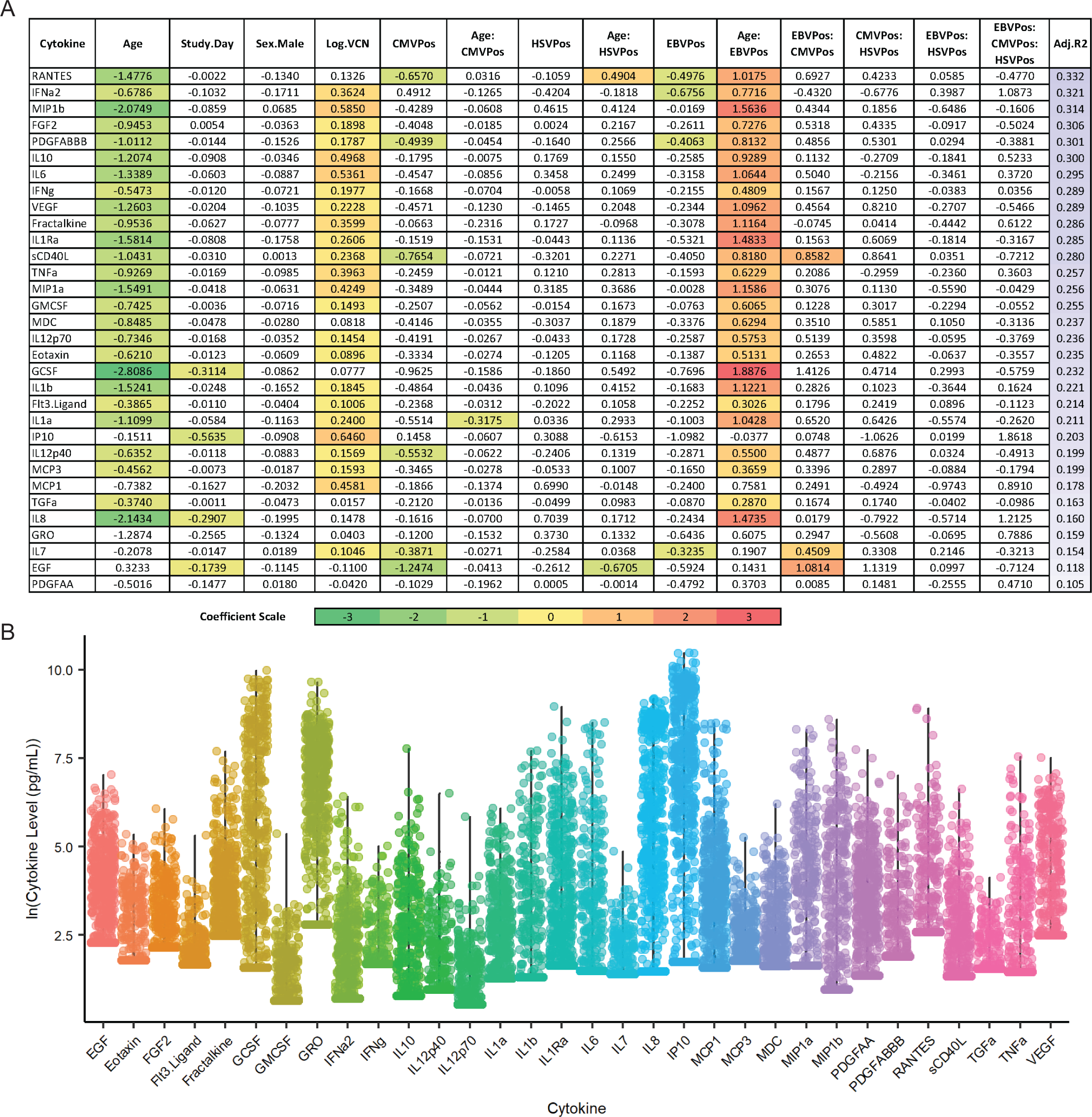
Heatmap summary of acute nasal wash cytokine regression models. Samples were from influenza infected subjects. (A) Summary of coefficient estimates for each factor included in the infected nasal wash linear regression model. Cytokine models are listed in decreasing order of the adjusted R^2^ value (shown on the far right) and coefficients are colored if FDR < 0.05. Red is associated with an increase in the cytokine level per unit change in the factor, and green is associated with a decrease in the cytokine level. (B) Acute nasal wash cytokine level distributions.

**Supplemental Table 1**. *Immune-related SNP genotype frequency in Memphis and Nicaragua*.

**Supplemental Table 2**. *Results from basal cell phenotype comparisons in PBMCs from healthy Nicaraguans and Memphians*.

**Supplemental Table 3**. *Gene expression Mann Whitney U and linear regression model results*.

**Supplemental Table 4**. *Regression results for eQTL analysis*. **Supplemental Table 5**. *Cytokine correlates of severity analysis results*. **Supplemental Table 6**. *Biological and herpesvirus severity associations*.

**Supplemental Table 7**. *Results from integrative analysis of biological, genetic, and herpesvirus effects on cytokine levels during influenza infection*.

## Acknowledgements

We thank the participants and clinical study support staff in each of the cohorts; Emily Walker and the Hartwell Center for processing Infinium Global Screening Array-24 v3.0 chips; Jessica Gaevert for technical assistance with RNA extractions; and ALSAC (American Lebanese Syrian Associated Charities). Select figures (Fig 1, Fig 3C, Fig 6E-F) were created using an academic license of BioRender software. This work was funded by: the Hercules Foundation – Belgium and the University of Antwerp (Methusalem funding; BOF Concerted Research Action), and the Research Foundation Flanders, project grant G.0409.12N and 1861219N (B.O.); the National Institute of Allergy and Infectious Diseases, US National Institutes of Health, under HHS contract HHSN272201400007C for Johns Hopkins Center of Excellence in Influenza Research and Surveillance (JH CEIRS) (A.P. and R.R.), HHS contract HHSN272201400006C for the St. Jude Center of Excellence for Influenza Research and Surveillance (SJ CEIRS) (R.W., S.S.C., P.G.T.), HHS contract 75N93021C00016 for the St. Jude Centers of Excellence for Influenza Research and Response (SJ CEIRR) (R.W., S.S.C., P.G.T.), HHS contract 75N93019C00052 for the Center for Influenza Vaccine Research for High Risk Populations (CIVR-HRP) of the Collaborative Influenza Vaccine Innovation Centers (R.W., S.S.C., P.G.T.), RO1 5R01AI120997 (A.G.), and RO1 AI107625 (P.G.T.).

The content is solely the responsibility of the authors and does not necessarily represent the official views of the National Institutes of Health.

## Author Contributions

Conceptualization, A.S. and P.G.T.; Methodology, A.S., E.K.A., L.T., and P.G.T.; Investigation, A.S., E.K.A., C.O., S.W., and T.J.; Formal Analysis, A.S., L.S., and S.P.; Visualization, A.S.; Writing – Original Draft, A.S.; Writing – Review & Editing, A.S., E.K.A., and P.G.T.; Resources, G.K., A.B., R.Z., K.S.S. J.C.D., S.H., G.E., P.V.D., V.V.T., B.O., R.W., S.S.C., A.P., R.R., A.G., P.G.T.; Supervision, P.G.T.; Funding Acquisition, B.O., R.W., S.S.C., A.P., R.R., A.G., P.G.T.

## Data Availability

Supporting data can be obtained from the authors upon reasonable request.

## Code Availability

Analyses were conducted using R, a publicly available software, as detailed in the Methods. All code is available upon request.

## Methods

### Study Subjects

#### Memphis Cohort

Plasma, nasal wash, and PBMCs were collected from healthy and influenza infected (n = 383) subjects in Memphis, Tennessee.^33^ Recruited subjects were 75% Black/African American and 25% White/Caucasian. To be included in the study, a flu positive patient must meet the clinical definition of influenza at the time of enrollment and symptoms must have begun within the previous 96 hours or less. Enrollment included index patients and household contacts (both flu positive and negative). Exclusion criteria included refusal to consent to nasal lavage samples, a history of receiving immunoglobulin or other blood products within the three months prior to enrollment, or any condition that in the opinion of the clinical site investigator place the subject at any risk or injury or render the subject unable to meet the study requirements. For index patients, nasal lavages and blood were collected at the day of enrollment (day 0) and days 3, 7, 10, and 28. Household contacts were also enrolled and provided nasal swabs on days 0. 3. 7, and 14, and 28, with blood on days 0 and 28. If at any point a household contact tested influenza positive, they were re-enrolled as a converted contact case and followed the same course of sample collection as the initial index patient. Symptom severity scores were collected throughout the duration of infection and were based on participant rankings using a visual analog scale and data consisted of five categories-total, upper respiratory, lower respiratory, gastrointestinal, and systemic. The peak total and lower respiratory symptom scores were utilized for severity analyses. Hospitalization status was also utilized and occurred in 25.9% of influenza positive participants. Written informed consent was obtained for participants or their guardians and written assent was acquired from age-appropriate subjects. This study was approved by the Institutional Review Boards of St. Jude Children’s Research Hospital and the University of Tennessee Health Science Center/Le Bonheur Children’s Hospital.

#### Colombia Cohorts

Healthy plasma samples were collected at a single time point from subjects in the local city of Calabazo (n = 100) and two indigenous populations, Seywiaka (n = 50) and Umandita (n = 50), which are 1.5 and 9.5 hours walk away from Calabazo, respectively.^35^ Recruited subjects were 100% Hispanic/Latino. All individuals willing to participate by completing the study questionnaires and permitting the withdrawal of blood and nasal swabs were included in the study. Exclusion criteria included children who are less than two years old when baseline enrollment is performed and anyone with a known immunosuppressive condition or ongoing receipt of immunosuppressive therapy. Signed consent or assent was obtained as appropriate and this study was approved by the Indigenous Health Council, the Tropical Health Foundation ethics committee, and the St. Jude Children’s Research Hospital Institutional Review Board.

#### Belgium Cohort

Healthy serum samples were collected at a single time point from 40 neonatal intensive care unit or pediatric unit nurses.^36^ Recruited subjects were 100% White/Caucasian. Inclusion criteria included a minimum of two years working in the respective hospital unit, a maximum of one year between blood sampling and last VZV exposure for pediatric unit nurses, and a minimum of one year between blood sampling and last VZV exposure for neonatal intensive care unit nurses. Exclusion criteria included immunosuppressed state (due to disease or medication), any malignancy in the last five years, receipt of blood products within three months of blood sampling, coagulation disorders, and any condition the investigator believed might interfere with the study. Written informed consent was obtained from all study participants and this study was approved by the ethics board of the Antwerp University Hospital, Antwerp, Belgium.

#### Baltimore and Taiwan Cohorts

Plasma and/or nasal wash samples were collected from healthy or influenza-infected subjects recruited from the East Baltimore and Bayview campuses in Baltimore, Maryland, USA (n = 225) and the Chang Gung Memorial Hospitals in the Taipei, Taiwan metropolitan area (n = 266).^37,38,62,63^ Recruited subjects from Taiwan were 100% Asian, and those from Baltimore were 67% Black/African American, 23% White/Caucasian, 1% American Indian or Alaskan Native, 9% Other. Possible index cases were recruited if, within the past 7 days of enrollment, they reported or documented fever and new or increased cough, headache, or sore throat. Cases were confirmed influenza positive if they received a positive influenza virus test result from the hospital visit (GeneXpert). Exclusion criteria included children younger than the age of five, currently incarcerated, unable to write or speak English, unable to provide informed consent, or unable to provide a telephone number for follow-up. Clinical coordinators obtained written informed consent, demographic, and clinical data. Samples were collected at the initial visit and at a 3-5 week follow-up visit. Additionally, symptom information included the presence or absence of wheezing and/or shortness of breath (WSOB), pneumonia (based on radiological or microscopic findings), hospitalization (for influenza associated illness), and administration of supplemental oxygen. In the Baltimore cohort, these symptoms occurred in 75%, 26%, 31%, and 17% of subjects, respectively. In the Taiwan cohort, the prevalence was 67%, 39%, 24%, and 19%, respectively. Due to the high prevalence of WSOB, it was excluded from downstream analyses for both cohorts. These studies were reviewed and approved by the Institutional Review Boards of Johns Hopkins School of Medicine and the Chang Gung Memorial Hospital (IRB00091667).

#### Nicaragua Cohort

Plasma samples were collected from healthy and influenza-infected subjects (n = 591) recruited at the Health Center Sócrates Flores Vivas (HCSFV) in Managua, Nicaragua.^34^ Recruited subjects were 100% Hispanic/Latino. Inclusion criteria include that an index case must live in the HCSFV catchment area, be positive for influenza by QuickVue Influenza A+B rapid test, experience onset of symptoms within the previous 48 hours, live with at least one additional person who agrees to participate in the study, and have no household contacts who have had influenza-like illness in the previous two weeks. There are no exclusion criteria for participation in this study. Household members were also recruited and monitored for influenza infection through 5 home visits, 2-3 days apart, over a 10-14 day period. Blood samples were collected at the initial visit and a 30 day follow-up visit, and participants submitted questionnaires regarding signs and symptoms of acute respiratory illness. Participants provided written informed consent and parental permission was obtained from parents or legal guardians of children, in addition to verbal assent from children aged six years and older. This study was approved by the Institutional Review Boards at the University of Michigan, Centro Nacional de Diagnóstico y Referencia (Ministry of Health, Nicaragua), and University of California, Berkeley.

### Herpesvirus Serostatus

Cytomegalovirus (CMV), Epstein-Barr virus (EBV), herpes simplex virus 1 (HSV1), herpes simplex virus 2 (HSV2), and human herpesvirus 6 (HHV6) serostatus was determined using plasma and the following ELISA kits, respectively: GWB-892399, GWB-DA817E, GWB-ECB3C0, GWB-C2349A, and Abnova KA1457.

### Cytokine Measurement

Plasma and nasal wash cytokine, chemokine, and growth factor levels were measured using the Luminex MAP system with the Milliplex HCYTOMAG-60K assay, and samples were processed according to the kit manual. Prior to statistical analyses, analytes were removed from the dataset if less than 15-40% of their values were above or below the lower and upper limits of detection, respectively. This value was selected based on total sample size and the prevalence of independent and dependent variables of interest.

### Flow Cytometry

PBMCs were stained for the following surface markers: Panel 1) CCR7-FITC (Biolegend Cat. 353216), CD25-APC (Biolegend Cat. 302610), CD3-PerCP eFluor710 (Thermo Cat. 46-0037-42), CD38-BV785 (Biolegend Cat. 303530), CD4-BV650 (BD Biosciences Cat. 563875), CD45RA-PE Dazzle/594 (Biolegend Cat. 304146), CD45RO-BV605 (Biolegend Cat. 304238), CD69-AF700 (BD Biosciences Cat. 310922), CD8-APC Fire750 (Biolegend Cat. 344746), gd TCR-PE/Cy7 (Biolegend Cat. 331222), PD1-BV711 (Biolegend Cat. 329928), TIGIT-PE (Biolegend Cat. 372704), and TIM3-BV421 (BD Biosciences Cat. 565562). Panel 2) CD11b-BV785 (Biolegend Cat. 301346), CD11c-PE/Cy7 (Biolegend Cat. 301608), CD123-BV421 (Biolegend Cat. 306018), CD14-APC eFluor780 (Thermo Cat. 47-0149-42), CD16-FITC (Biolegend Cat. 360715), CD19-PerCP eFluor710 (Thermo Cat. 46-0198-42), CD1c-BV711 (Biolegend Cat. 331536), CD3-PerCP eFluor710 (Thermo Cat. 46-0037-42), CD56-BV605 (Biolegend Cat. 318334), CD68-APC (Biolegend Cat. 333810), CD80-PE (Biolegend Cat. 305208), and HLA DR-BV650 (Biolegend Cat. 307650). Panel 3) CD3-AF700 (Biolegend Cat. 300323), CD4-APC Fire750 (Biolegend Cat. 300559), FOXP3-PE (Biolegend Cat. 320107). Briefly, after thawing from cryopreservation and plating in a 96-well round bottom plate, cells were spun down and resuspended in 50μL of human Fc block (BD Biosciences Cat. 564220) in DPBS and incubated for 10 minutes at room temperature. Afterwards, 50μL of a Live/Dead Aqua (Tonbo Cat. 13-0870-T100) and pre-titrated surface antibody cocktail in DPBS were added to each well and cells were incubated for 30 minutes on ice and in the dark. Additionally, the BD Cytofix/Cytoperm kit (BD Biosciences Cat. 554714) and FOXP3 Transcription Factor Staining Buffer set (eBioscience Cat. 00-5523-00) were used for intracellular staining of CD68-APC (Panel 2) and FOXP3-PE (Panel 3), respectively and in accordance with the kit manual. Cells were washed, resuspended in FACS (fluorescence-activated cell sorting) buffer and analyzed on a BD Fortessa cell analyzer and analyzed using FlowJo software.

### Microneutralization Assays

Microneutralization titers were performed according to guidelines outlined by the World Health Organization (WHO) (Global Influenza Surveillance Network, 2011). Reported results are the reciprocal dilution which inhibited virus growth as determined by an ELISA readout or hemagglutination assay.

### Viral Titers

Influenza viral load was measured with quantitative reverse-transcription real-time polymerase chain reaction (qRT-PCR) assays and RNA from nasal swabs. Nucleic acid was extracted from samples using the MagMAXtm-96 AI/ND Viral RNA Isolation kit on the KingFisher magnetic processor platform. Afterwards, a 25mL qRT-PCR reaction was set up utilizing the Qiagen One-Step RT-PCR kit, influenza A matrix or influenza B non-structural protein primers and the protocol outlined by the WHO(Global Influenza Surveillance Network, 2011).

### Human Genetic Studies

#### Prioritization of SNPs

The UCSC Human Genome Browser build GRCh37/19 was used to curate a list of 308 common SNPs in genes encoding 41 cytokines measured by the Milliplex assay HCYTOMAG-60K and for 3 transcription factors important in initiating antiviral immunity, namely IRF3, IRF7, and all 3 subunits of NFkB. Briefly, putative functional SNPs were prioritized and included the following classes of SNPs: missense, splice, non-coding, and those which are located in the 3’ or 5’ untranslated region (UTR) of the promoter region. Utilizing 1000 Genomes Project data, the curated list of SNPs was reduced to 209 by filtering for SNPs with minor allele frequencies (MAF) which significantly vary across ancestrally distinct populations. For a given SNP, this was calculated using the MAFs of the African (AFR), American (AMR), Eastern Asian (EAS), European (EUR), and South Asian (SAS) populations. A SNP was considered to significantly vary if the standard deviation of the MAFs across these populations was greater than 5. Lastly, the prioritized list of SNPs was reduced to 94 following a tagSNP analysis to identify the most informative SNPs in a region based on linkage disequilibrium. The tagSNP analysis was conducted by utilizing the SNPclip tool from the National Cancer Institute (https://ldlink.nci.nih.gov/?tab=snpclip) and data from African (YRI), African American (ASW), Mexican (MXL), and European (CEU) populations, as these most reflect the populations in this study.

#### DNA Extraction

For the Memphis cohort, 90% of samples were extracted in a clinical HLA core at St. Jude, the remaining 10% were extracted from peripheral blood mononuclear cells using the Zymo Quick-DNA Microprep Plus Kit (Cat. D4074). All DNA samples for the Nicaragua cohort were extracted from 200mL of neutrophil pellets using the Qiagen QIAamp DNA Mini Kit (Cat. 51306). All samples were quantified using a Nanodrop, the average 260/280 ratio was 1.86 (range 1.16 - 2.04). For Fluidigm SNP Type assays, all samples met the minimum DNA input requirement of 10ng/mL, and all ancestry informative marker samples had an input of 250ng.

#### SNP Genotyping

Genotypes for the curated list of 94 SNPs were determined using the Fluidigm SNP Type Assay on the BioMark HD System and 96.96 Dynamic Array integrated fluidic circuits. SNP Type assays were designed using Fluidigm’s D3 Assay Design Tool. The assay was performed in accordance with the Fluidigm protocol, using PCR grade water as a negative control, and including the specific target amplification step with 14 cycles of amplification.

#### Ancestry Informative Markers

Genetic based ancestry was determined using the Infinium Global Screening Array-24 v3.0 BeadChip which measures 654,027 markers including multiethnic genome-wide content, curated clinical research variants, and quality control markers. High quality DNA was submitted to the St. Jude Hartwell Center for preparation and processing.

### Gene Expression Assays

#### PBMC Stimulation

PBMCs from 188 subjects (94 Memphians and 94 Nicaraguans) were plated at a max of 850,000 cells per well and resuspended in 1μg/mL of lipopolysaccharide, Pam3CSK4, transfected poly(I:C), transfected poly(dA:dT), or plain media as an unstimulated control. Plates were incubated at 37°C for 24 hours. Following stimulation, plates were spun at 500g for 5 minutes. Supernatants were transferred to clean 96-well plates and stored at −80°C. Pellets were resuspended in 300μL of lysis buffer and stored at −80°C in a labeled epitube until ready for extraction.

#### Gene Expression

RNA was extracted using the Zymo Quick-RNA extraction kit (Cat. R1050), and stored in a 96-well plate at −80°C until ready for PCR. Prior to running gene expression assays, stored RNA was thawed and normalized to the highest amount of RNA possible, with respect to stimulation condition. The input RNA was 14ng for unstimulated control, 18ng for Pam3CSK4, 40ng for lipopolysaccharide, 50ng for transfected poly(I:C), and 50ng for transfected poly(dA:dT). The expression of 96 immune genes was measured using the Fluidigm Fast Gene Expression Analysis Using EvaGreen on the BioMark HD System and 96.96 Dynamic Array integrated fluidic circuits. Primers were designed using Fluidigm’s D3 Assay Design Tool. The assay was performed in accordance with the Fluidigm protocol, including the specific target amplification step with 16 cycles of amplification, and 2 water samples per chip as negative controls. Quality control of gene expression results included removing samples/genes with no measurable gene expression or samples with melting curve peaks that were more than one degree away from the consensus peak melting temperature. Following QC steps, 4 genes were removed altogether (*IFNL2, IL9, SFTPD, TMEM173*) and 92 remained for further analysis (2 housekeeping - *ACTB, GAPDH*; 90 immune - *AIM2, AREG, BIK, CCL11, CCL2, CCL22, CCL3, CCL4, CCL5, CCL7, CD40LG, CD69, CD80, CD86, CGAS, CSF3, CX3CL1, CX3CR1, CXCL1, CXCL10, CXCL2, DDX58, EGF, EOMES, FGF2, FLT3LG, GATA3, GZMB, IFI16, IFIT1, IFITM3, IFNA2, IFNB1, IFNG, IFNL1, IL10, IL12A, IL12B, IL13, IL15, IL17A, IL18, IL1A, IL1B, IL1RN, IL2, IL4, IL5, IL6, IL8, IRF1, IRF3, IRF7, IRF9, ISG15, JUN, KLRC1, KLRK1, LTA, MAVS, MX1, MX2, NFATC1, NFKB1, NFKB2, NFKBIA, NLRP3, OAS1, PDGFA, PDGFB, PTGS2, REL, RELA, RELB, RORC, SELL, SOCS1, SOCS3, STAT1, STAT3, STAT4, TBK1, TBX21, TGFA, TGFB1, TNF, TNFAIP3, TREX1, VEGFA, VEGFC*). Lastly, prior to conducting comparative analyses, results were normalized again using housekeeping gene expression (*ACTB*).

### Luciferase Assays

Determination of IL10-rs3024496 and IRF3-rs2304204 as functional SNPs was achieved using luciferase assay constructs ordered from GenScript. For IL10-rs3024496, a 1,1114bp region (chr1:206,945,768-206,946,881, GRCh37/19), encompassing the IL10 promoter, was inserted upstream of the *luc2* gene in the pGL4.10[*luc2*] plasmid (Promega Cat. E6651). Following this step, two final constructs were created, one with the C allele and one with the T allele for rs3024496, by cloning a 2,364bp region (chr. 1:206,939,683-206,942,046, GRCh37/19) that encompassed the IL10 3’UTR and the SNP of interest. For IRF3-rs2304204, a 1,394bp region (chr19:50,168,220-50,169,613, GRCh37/19), encompassing the IRF3 promoter, was inserted upstream of the *luc2* gene in the pGL4.10[*luc2*] plasmid (Promega Cat. E6651). As the SNP was located within the promoter region, two versions of this construct were created and cloned, one with the G allele and one with the A allele for rs2304204.

A549s (ATCC Cat. CCL-185) were transfected with each plasmid using the TransIT LT Transfection Reagent (Mirius Bio Cat. MIR 2300). Briefly, 1.5×10^5^ cells/well were seeded into 12-well plates, and incubated overnight at 37°C and 5% CO_2_. The following day, 1μg of pGL4.10 test plasmid and 0.01μg of pRL-TK control plasmid (Promega Cat. E2241) were added per well, according to the TransIT LT protocol. All transfections were conducted in triplicate. At 24 hours post transfection, cells were collected and luciferase expression was measured using the Dual-Luciferase Assay kit (Promega Cat. E1910) on the GloMax platform in accordance with the kit protocol. For influenza stimulated results, at 24 hours post transfection, cells were washed with PBS, 1mL of 6.0×10^5^ pfu/mL Influenza A virus (A/Singappore/INFIMH-16-019/2016, multiplicity of infection = 4) was added to each well, and plates were incubated at 37°C and 5% CO_2_ for 2 hours. Untreated control plates received infection media instead of inoculum. Following the 2 hour incubation, inoculum was removed, cells were washed with PBS, 1mL of infection media was added to each well, and plates were incubated for an additional 24 hours at 37°C and 5% CO_2_, after which luciferase expression was measured.

### Statistical Analysis

Detailed descriptions of specific applied statistical approaches can be found in figure legends and/or the main text. All analyses were performed with R using base functions (lm, p.adjust) and generally the following packages: Rtsne, GGally, pheatmap, ggpubr, broom, formattable, and tidyverse.^64–70^ Dimension reduction techniques included principal component analysis and t-Distributed Stochastic Neighbor Embedding. For the cytokine tSNE analysis, any missing values were replaced with the median of that cytokine level, with respect to the cohort. Pearson and Spearman correlations were used to determine cytokine correlates of severity for continuous symptom outcomes. The unstimulated gene expression PCA required complete observations (i.e. no missing values), and thus was generated from 102 subjects and 67 genes. Multivariate analyses were conducted utilizing logistic or linear regression modeling, depending on whether the outcome variable was binary or continuous, respectively. For all models, continuous variables were standardized using the scale() function in R. All remaining analyses were Mann-Whitney U or Kruskal Wallis tests, depending on whether there were two or more comparison groups, respectively. All herpesvirus association analyses only included CMV, EBV, and HSV, due to the overall high seroprevalence of HHV6 (%) and VZV (%). When appropriate and indicated, p-values were adjusted for multiple comparisons using the Benjamini-Hochberg method for controlling the false discovery rate.^71^

